# A geostatistical approach to estimate high resolution nocturnal bird migration densities from a weather radar network

**DOI:** 10.1101/690065

**Authors:** Raphaël Nussbaumer, Lionel Benoit, Grégoire Mariethoz, Felix Liechti, Silke Bauer, Baptiste Schmid

## Abstract

1. Quantifying nocturnal bird migration at high resolution is essential for (1) understanding the phenology of migration and its drivers, (2) identifying critical spatio-temporal protection zones for migratory birds, and (3) assessing the risk of collision with man-made structures.
2. We propose a tailored geostatistical model to interpolate migration intensity monitored by a network of weather radars. The model is applied to data collected in autumn 2016 from 69 European weather radars. To cross-validate the model, we compared our results with independent measurements of two bird radars.
3. Our model estimated bird densities at high resolution (0.2°latitude-longitude, 15min) and assessed the associated uncertainty. Within the area covered by the radar network, we estimated that around 120 million birds were simultaneously in flight [10-90 quantiles: 107-134]. Local estimations can be easily visualized and retrieved from a dedicated interactive website: birdmigrationmap.vogelwarte.ch.
4. This proof-of-concept study demonstrates that a network of weather radar is able to quantify bird migration at high resolution and accuracy. The model presented has the ability to monitor population of migratory birds at scales ranging from regional to continental in space and daily to yearly in time. Near-real-time estimation should soon be possible with an update of the infrastructure and processing software.

## Introduction

Every year, several billion birds undergo migratory journeys between their breeding and non-breeding grounds (Dokter et al., 2018; Hahn, Bauer, & Liechti, 2009). These migratory movements link ecosystems and biodiversity at a global scale (Bauer & Hoye, 2014), and their understanding and protection requires international efforts (Runge, Martin, Possingham, Willis, & Fuller, 2014). Indeed, declines in many migratory bird populations (Sanderson, Donald, Pain, Burfield, & van Bommel, 2006; Vickery et al., 2013) resulted from the rapid changes of their habitats, including the aerosphere (Diehl, 2013). Changes in aerial habitats are diverse, and their consequences still poorly resolved. Nevertheless, climate change may alter global wind patterns and consequently the wind assistance provided to migrants (La Sorte, Horton, Nilsson, & Dokter, 2019). Likely more severe, the impact of direct anthropogenic changes include for instance light pollution that reroute migrants (Van Doren et al., 2017), or buildings (Winger et al., 2019), wind energy production (Aschwanden et al., 2018), and aviation (van Gasteren et al., 2019) that together cause billions of fatalities every year (Loss, Will, & Marra, 2015).

In the face of these threats and for setting up efficient management actions, we need to quantify bird migration at various spatial and temporal scales. Fine scale monitoring is crucial for understanding the phenology of migration and its drivers, identifying critical spatio-temporal protection zones to enhance conservation actions, and assessing collision risks with human-made structures and aviation to inform stakeholders. However, the great majority of migratory landbirds fly at night (Winkler, 1999), challenging the quantification of the sheer scale of bird migration.

Radar monitoring has the potential to quantify migratory movements of birds at the continental scale (Drake & Bruderer, 2017). Initially limited to single dedicated short-range measurements, the use of existing weather radar networks provide continuous monitoring over large geographical areas such as Europe or North America (Gauthreaux, Belser, & van Blaricom, 2003; Shamoun-Baranes et al., 2014), and led to an upswing of radar aeroecology (Bauer et al., 2017; Dokter et al., 2018; Van Doren & Horton, 2018). One important challenge in using networks of weather radars is the interpolation of their signals in space and time. Recent studies (Dokter et al., 2018; Van Doren & Horton, 2018) have used relatively simple interpolation methods as they targeted patterns at coarse spatial and/or temporal scales. However, these methods are insufficient if higher spatial or temporal resolution is wanted such as for the fundamental and applied challenges outlined above.

To achieve high resolution interpolation of migration intensity derived from weather radars (20km-15min), we propose a tailored geostatistical framework able to model the spatio-temporal pattern of bird migration. Starting from time series of bird densities measured by a radar network, our geostatistical model produces a continuous map of bird densities over time and space. A major strength of this method is its ability to provide the full range of uncertainty and thus, to evaluate the probability that bird densities reaches a given threshold. In addition to the estimation map, the method also produces simulation maps which are essential for several applications such as quantification of the total number of birds.

As a proof of concept, we applied our geostatistical model to a three weeks dataset from the European Network of weather radar (Huuskonen, Saltikoff, & Holleman, 2014) and validated the results with independent dedicated bird radar data. In addition to insights into the spatio-temporal scales of broad front migration, our approach provides high resolution (0.2°latitude and longitude, 15min) interactive maps of the densities of migratory birds.

## Materials and Methods

### Weather radar dataset

Our dataset originates from measurements of 69 European weather radars, spread from Finland to the Pyrenees (8 countries) and covering the period from 19 September to 10 October 2016 (Figure 1). It thus encompasses a large part of the Western-European flyway during fall migration 2016.

**Figure 1.**
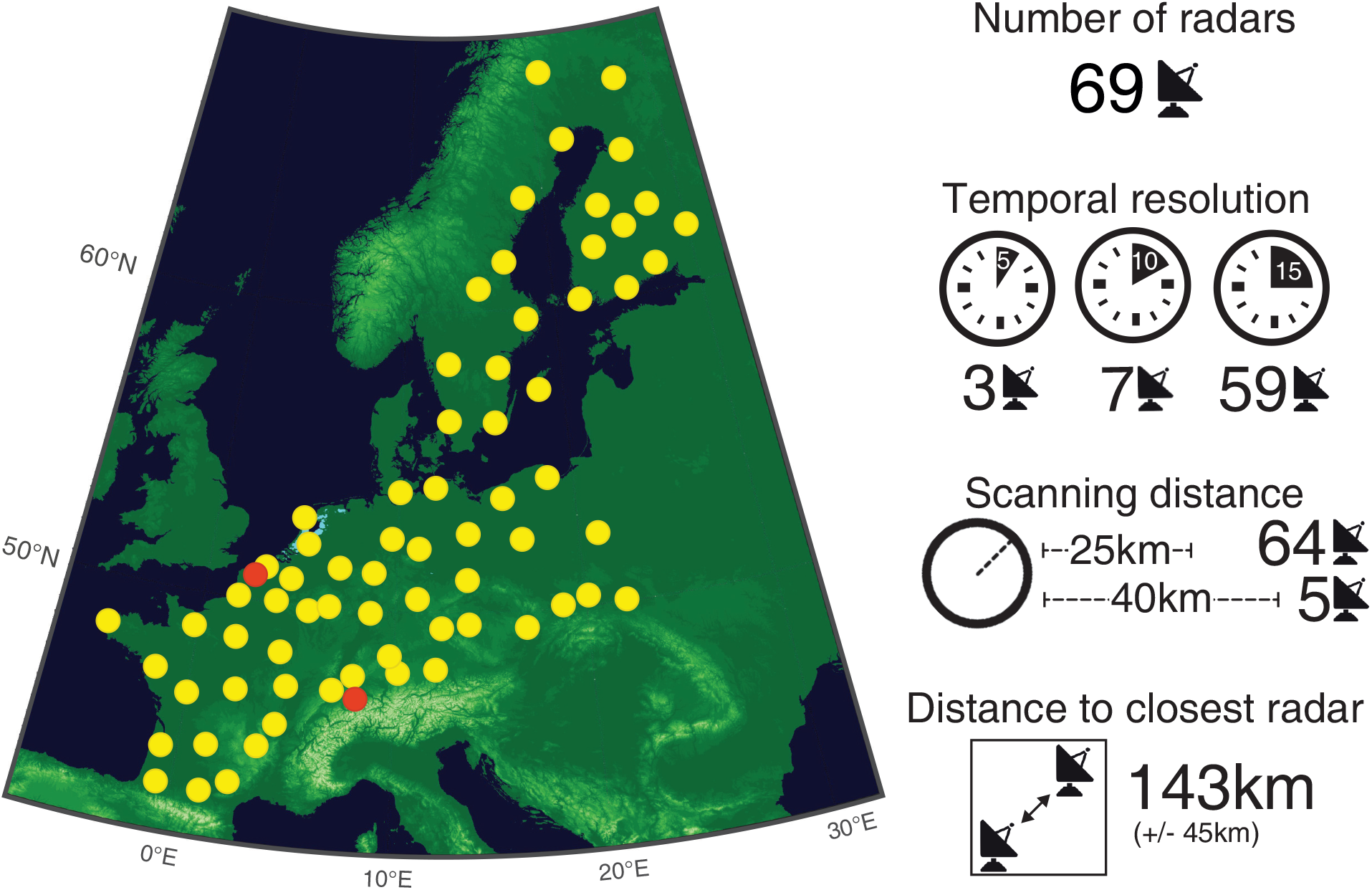
(Left) Locations of weather radars of the ENRAM network, whose fall 2016-data were used in this study (yellow dots), and their key characteristics (right panel). We used data from two dedicated bird radars – in Switzerland and France - for validation (red dots).

Based on the reflectivity measurements of these weather radars, we used the bird densities as calculated and stored on the repository of the European Network for the Radar surveillance of Animal Movement (ENRAM) (github.com/enram/data-repository)(see (Nilsson et al., 2019) for details on the conversion procedure). We inspected the vertical profiles and manually cleaned the bird densities data (see detailed procedure in Supporting Information S1).

Since we targeted a 2D model, we vertically integrated the cleaned bird densities from the radar elevation and up to 5000 m above sea level. Because we aimed at quantifying nocturnal migration, we restricted our data to night time, between local dusk and dawn (civil twilight, sun 6°below horizon). Furthermore, as rain might contaminate and distort the bird densities calculated from radar data, a mask for rain was created when the total column of rain water exceeds a threshold of 1mm/h (ERA5 dataset from (Copernicus Climate Change Service (C3S), 2017). In the end, the resulting dataset consisted of a time series of nocturnal bird densities [bird/km^2^] at each radar site with a resolution of 5 to 15 min (Figure 2).

**Figure 2.**
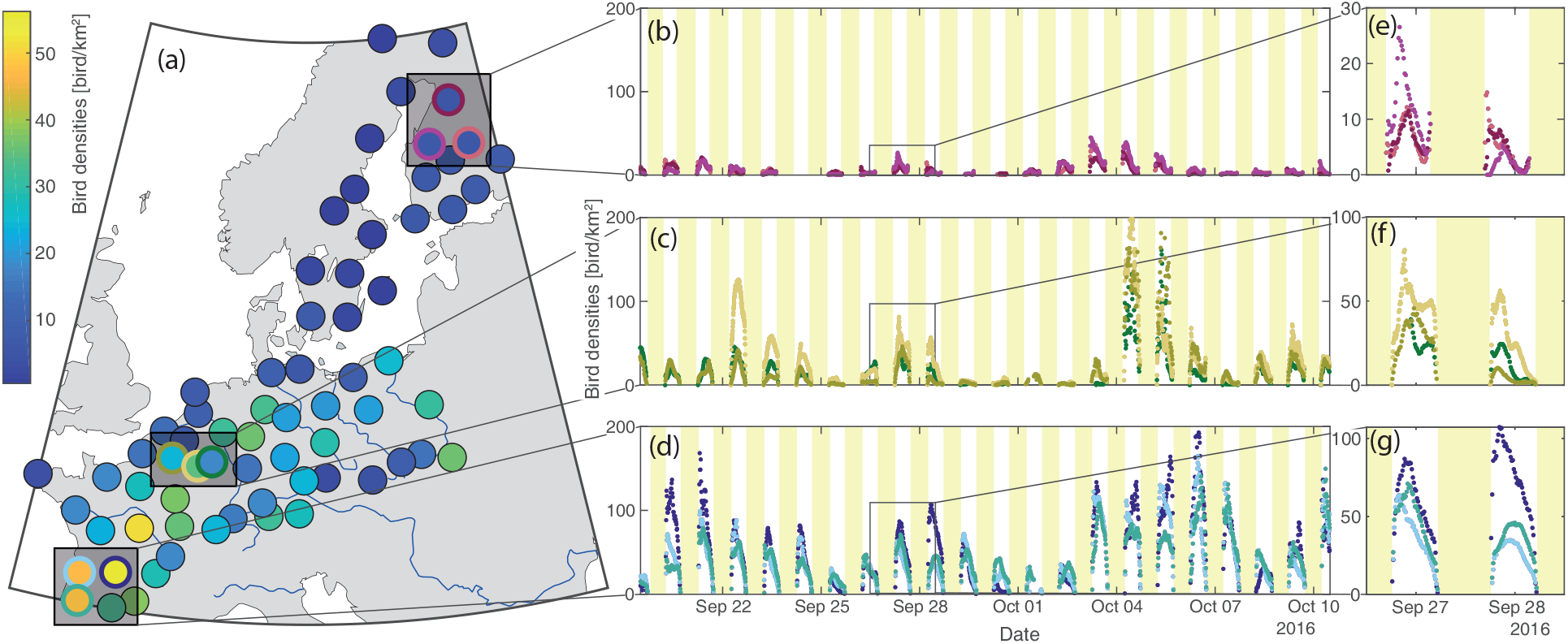
Space-time variability of bird densities as measured by a radar network. (a) Average bird densities over the whole study period (time series of radar with a coloured outer circle represented in the subsequent panels, respectively). (b-d) Time series of bird densities measured at different locations (colour of the dots corresponds to the colour of the outer circle in panel a). (e-g) Zoom on a two-days period.

### Interpolation approach

Bird densities are strongly correlated both spatially at continental (Figure 2a-d) to regional scales (Figure 2c), and temporally at daily (Figure 2b-d) to sub-nightly scales (Figure 2e-g).

These strong spatio-temporal correlations motivated the choice of using a geostatistical framework to interpolate the punctual radar observations. In such framework, bird densities are considered as a space-time random process that is fully defined by its covariance matrix (e.g. Chilés & Delfiner, 1999). To perform optimally, however, geostatistical interpolation requires bird densities to be stationary (i.e. mean, variance and covariance) in both space and time (e.g. Chilés & Delfiner, 1999) – an assumption hardly ever satisfied for migratory events. Rather, there are two main and obvious non-stationarities in our dataset: (1) migration is more intense in the south than in the north of Europe (Figure 2a), and (2) migration is more intense in the middle of the night than during twilight (Figure 2e-g). To account for these non-stationarities, we develop a tailored geostatistical model that decomposes the migration signal into four independent components.

### Geostatistical model

The bird density *Z*(**s***, t*) observed at location **s** and at time *t* is modelled by

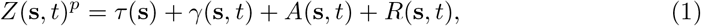

where *p* is a power transformation, *τ* the continental trend, *c* is the curve describing the nightly trend, *A* the nightly amplitude, and *R* the residual term (Figure 3). A power transformation is used on bird densities in order to transform the highly skewed marginal distribution into a Gaussian distribution (Figure S3-1 in Supporting Information S3). The trend and the curve are deterministic functions accounting for the two non-stationarities, whereas the amplitude and the residuals are stationary random processes modelling the spatio-temporal variability at nightly and sub-nightly scales respectively. The four components of the model are detailed in the following sub-sections.

### Trend

The increasing bird densities southwards (Figure 2a) create a first non-stationary in the dataset. Although this trend changes over the year, it can be considered constant over the short study period. Therefore, we model the trend as a deterministic planar function

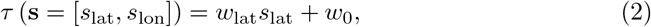

where *s*_lat_ and *s*_lon_ are latitude and longitude of location **s**, *w*_lat_ is the slope coefficient in latitude and *w*_0_ is the value of the trend at the origin. Because no longitudinal trend is observed in the data (Figure 1a), only latitude is used to parametrize the trend function (see Figure S3-2 in Supporting Information S3). It is worth noting that if longer periods are considered, Eqn. 2 should be replaced by a more complex parametric function in order to handle the emerging patterns of long term non-stationarity.

**Figure 3.**
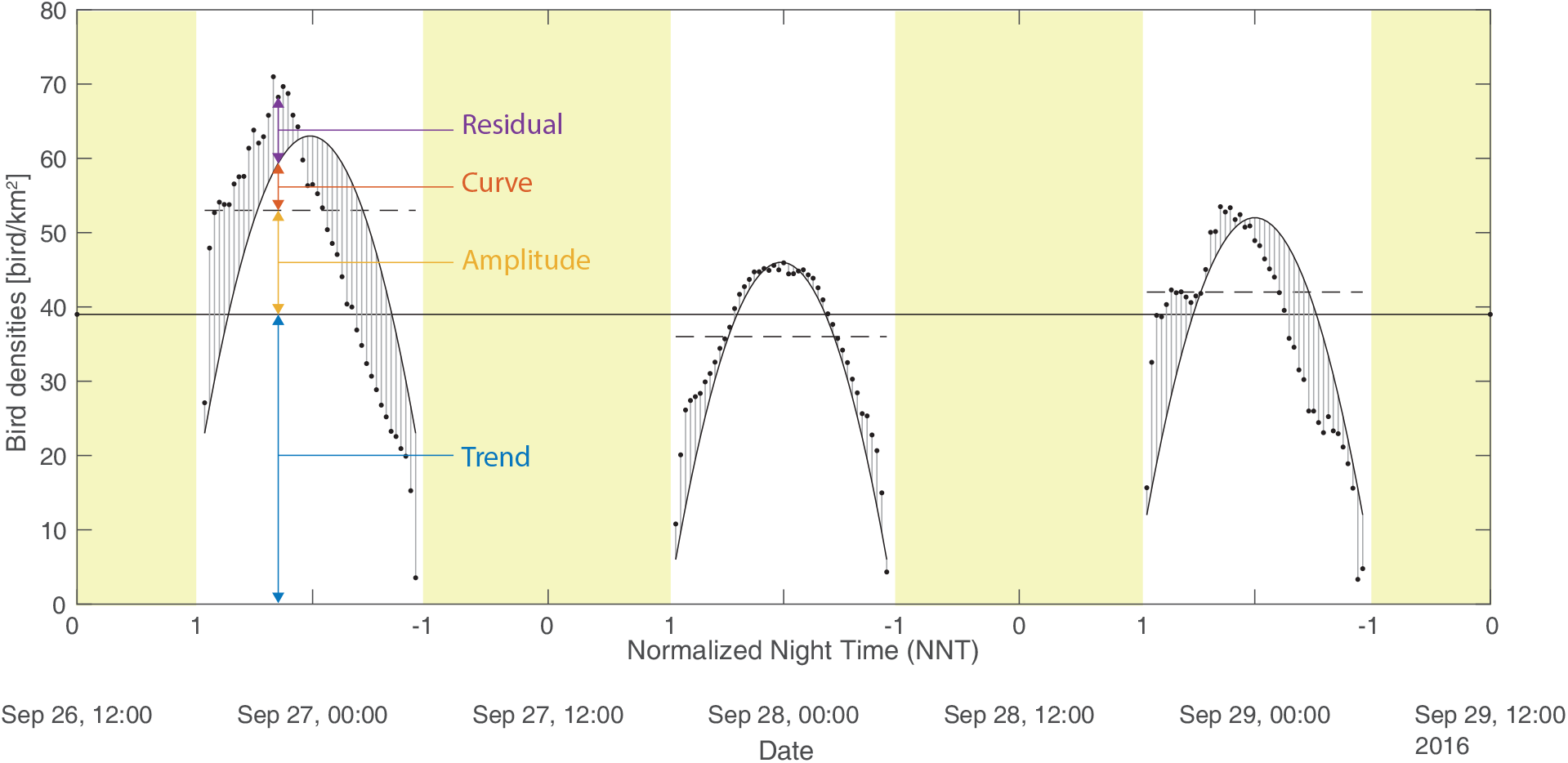
Illustration of the proposed mathematical model decomposition into trend, amplitude, curve and residual. Note that the power transformation was not applied on this illustration.

### Curve

The second non-stationarity visible in the dataset is the nightly pattern (Figure 2e-g) that results from the onset and sharp increase of migration activity after sunset, and its slow decrease towards sunrise (e.g. Bruderer & Liechti, 1995). Similar to the trend, this pattern needs to be extracted from the original signal to avoid non-stationarity. This is done using a curve template *c* for all nights and locations, defined as the polynomial of degree 6

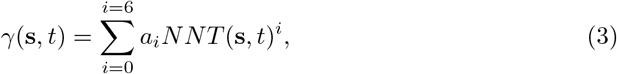

where *a*_*i*_ are the coefficients of the polynomial and *N N T* (Normalized Night Time) is the standardized proxy of the progression of night defined as

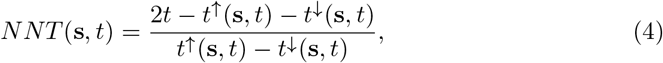

where *t*^↓^(**s**, *t*) and *t*^↑^(**s**, *t*) are the times of civil dusk and dawn respectively. *N N T* is defined such that the local sunrise or sunset occur at *N N T* = *−*1 and *N N T* = 1, respectively.

### Amplitude

After removing the non-stationarities with the trend and the curve, the amplitude *A* models the nocturnal bird densities at the daily scale as a stationary space-time random process. Its value is therefore constant within a night at a given location but varies between locations and between nights. It accounts for the correlation at the scale of several hundred kilometers and several days (Figure 1).

### Residual

The variation in bird densities not modeled by trend, curve and amplitude, is still strongly correlated in space and time at the hourly scale (Figure 2e-g).

### Model parameterization

The values of the model parameters are determined by fitting the model to the observed bird densities. In the Supporting Information S3, the parametrisation procedure is detailed and the significance of the resulting parameters of the model are discussed.

### Bird migration mapping

After parametrisation, the geostatistical model can be used to interpolate bird density observations derived from weather radars to produce high resolution maps. The full mathematical description of the procedure is detailed in Supporting Information S2 and a brief outline is given below.

The estimation of the bird density at any unsampled location *Z*(**s**_0_*, t*_0_)* is performed by first estimating independently each component, and then recombining them according to Eqn. 1. Estimating the deterministic components (i.e. trend and curve) is straightforward since they can be computed by applying Eqn. 2 and 3 at the target location and time **s**_0_*, t*_0_. Since the probabilistic components (amplitude and residual) are modelled as random processes, the estimations of *A*(**s**_0_*, t*_0_)* and *R*(**s**_0_*, t*_0_)* are performed by kriging (e.g. Chilés & Delfiner, 1999; Goovaerts, 1997). An important advantage of using kriging is that it expresses the estimation as a Gaussian distribution, thus providing not only the “most likely value” (i.e. mean or expected value) but also a measure of uncertainty with the variance of estimation. Tracking back the uncertainty provided by kriging to the final estimation *Z*(**s**_0_*, t*_0_)* is non-trivial but possible through the use of a quantile function (see Supporting Information S2 for details). Consequently, the estimation *Z*(**s**_0_*, t*_0_)* is expressed as the median and its uncertainty range is defined as the quantiles 10 and 90. A continuous space-time estimation (with uncertainty) map is then computed by repeating the procedure for estimating a spatio-temporal point **s**_0_*, t*_0_ on a discrete set of points (i.e. grid).

In addition to the kriging estimation, we also provide simulation maps. Although kriging is known to produce accurate point estimates, it leads to excessively smooth interpolation maps (e.g. Goovaerts, 1997) and thus fails to reproduce the fine-scale texture of the process at hand. Consequently, estimation should be complemented by geostatistical simulation (e.g. Chilés & Delfiner, 1999; Goovaerts, 1997) in applications for which the space-time structure of the interpolation map matters (e.g. when non-linear transformations are applied to the interpolated bird densities map, or when aggregation in space or time is required). However, simulations come with a heavy computational cost as a large ensemble of realisations is required to quantify the uncertainty associated with the interpolation.

In the case study presented in this paper, both estimation and simulations maps are calculated on a spatio-temporal grid with a resolution of 0.2°in latitude (43°to 68°) and longitude (−5°to 30°) and 15 minutes in time, resulting in 127×176×2017 nodes. Over this large data cube, the estimation and simulation are only computed at the nodes located (1) over land, (2) within 200km of the nearest radar and (3) during nighttime (*N N T* (*t*, **s**) *< −*1 or 1 *< N N T* (*t*, **s**)).

## Validation

### Cross validation

We tested the internal consistency of the model by cross-validation. It consists of sequentially omitting the data of a single radar, then estimating bird densities at this radar location with the model and finally, comparing the model-estimated value and *Z*(**s***, t*)* observed data *Z*(**s***, t*). The model is assessed by its ability to provide both the smallest misfit errors, i.e. ||*Z*(**s***, t*)* − Z(**s***, t*)||, and uncertainty ranges matching the magnitude of these errors. Because it is difficult to quantify both aspects for a non-normalized variable, the normalized error of estimation is used on the power transformation variable

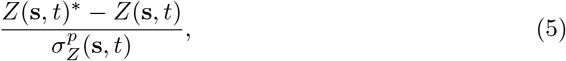

where 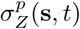 is the standard deviation of the estimation as defined in Eqn. S2-10 of Supporting Information S2.

### Comparison with dedicated bird radars

A second validation of our modelling framework (from data acquisition by weather radars to geostatistical interpolation) is to compare model-predicted bird migration intensities with the measurements of two dedicated bird radars (Swiss BirdRadar Solution AG, swiss-birdradar.com) located in Herzeele, France (50°53’05.6”N 2°32’40.9”E) and Sempach, Switzerland (47°07’41.0”N 8°11’32.5”E). These bird radars register single echoes transiting through the radar beam, allowing to compute migration traffic rates (MTR) and average speed of birds aloft (Nilsson et al., 2018; Schmid et al., 2019).

## Results

### Validation

#### Cross validation

In our study, the normalized error of estimation over all radars has a near-Gaussian distribution with mean 0.017 and variance of 0.60 (Figure S4-1 in Supporting Information S4). The near-zero mean of the error distribution indicates that our model provides non-biased estimations of bird densities, while the near-one standard deviation supports that the model provides appropriate uncertainty estimates, albeit slightly overconfident. The performance of the cross-validation shows radar-specific biases (i.e. constant under- or over-predictions)(Figure S4-2 in Supporting Information S4). The biases are not spatially correlated (Figure S4-3 in Supporting Information S4) and therefore such biases do not originate from the geostatistical model itself (or of country-specific data quality). Rather, these radar-specific biases probably come from either the data, such as birds non-accounted for (e.g. flying below the radar), or error in the cleaning procedure (e.g. ground scattering). In contrast, the variances of the normalized error of estimation of each radar are close to 1 and thus, demonstrate the accuracy of the estimated uncertainty range (Figure S4-2 and Figure S4-3 in Supporting Information S4).

#### Comparison with dedicated bird radars

The daily migration patterns estimated by our model coincide generally well with the observations derived from dedicated bird radars (Figure 4). First, the estimations of the model correctly reproduce the night-to-night migration intensity, with the exception of a few nights (27-30 September for Herzeele and 26/27 September for Sempach). Second, the intra-night fluctuations are also properly reproduced (e.g. 4/5 October for both radars). Over the whole validation period, the normalized estimation error has a mean of 0.6 and a variance of 1.3 at Herzeele radar location (n=164), and a mean of −0.8 and a variance of 1.3 at Sempach radar location (n=264). These normalized estimation errors indicate a tendency of the model to slightly overestimate bird densities in Herzeele and to underestimate it in Sempach. Finally, both variances were close to one, which shows that the model provides a reliable uncertainty range. Overall, the root-mean-square error of the non-transformed variable (i.e. the actual bird densities) was around 20 bird/km^2^ for both radars, which demonstrates the good performance of the model at these two test locations.

**Figure 4.**
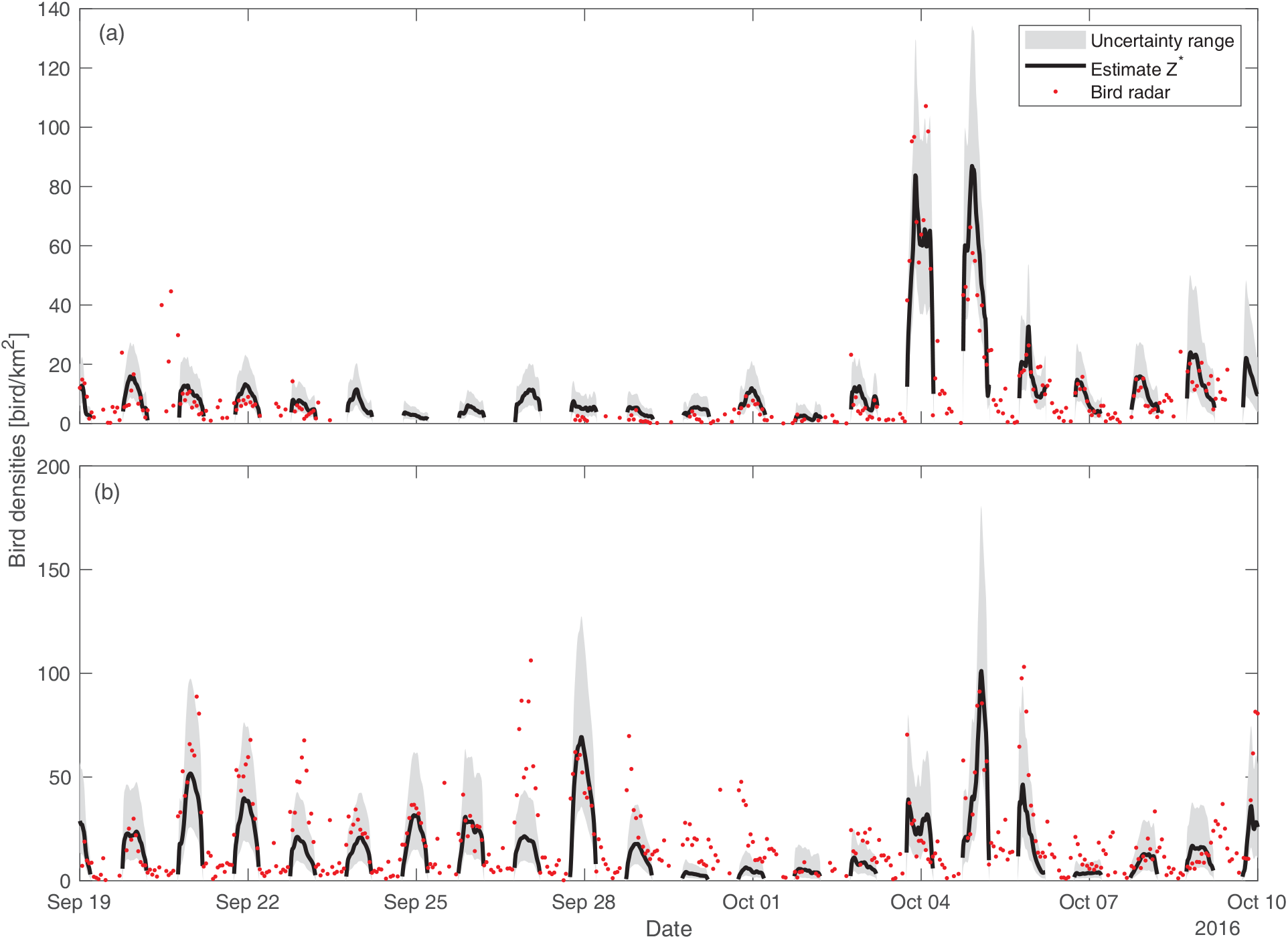
Comparison of the estimated bird densities (black line, 10-90 quantiles uncertainty range in grey) and the bird densities (red dots) observed using dedicated bird radars at two locations in (a) Herzeele, France (50°53’05.6”N 2°32’40.9”E) and (b) Sempach, Switzerland (47°07’41.0”N 8°11’32.5”E). Note that because of the power transformation, model uncertainties are larger when the migration intensity is high. It is therefore critical to account for the uncertainty ranges (light gray) when comparing the interpolation results with the bird radars observations (red dots).

#### Application to bird migration mapping

The main outcome of our model is to estimate bird densities at any time and location within the domain of interest. This is illustrated by the estimation of bird densities time series at specific locations (e.g. Figure 4), and by the generation of bird densities maps at different time steps (Figure 5).

**Figure 5.**
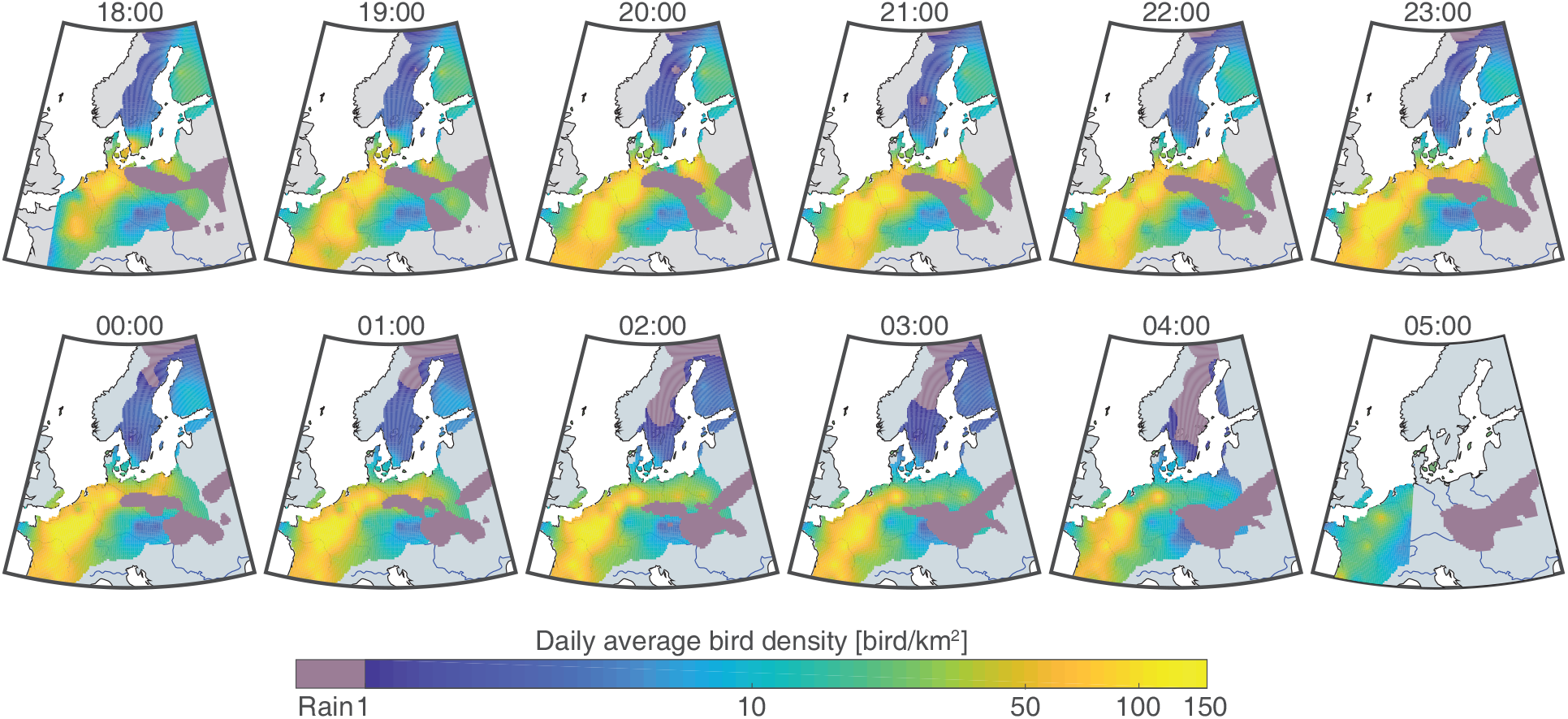
Maps of bird densities estimation every hour of a single night (4/5 October). Civil sunset and sunrise limits are visible on the first and last snapshots. The highest bird densities are in the corridor from Northern Germany to southwestern France. Rain limits migration in Southern Poland, Czech Republic and Southern Germany. The sunrise and sunset fronts are visible at 18:00 and 05:00 with lower densities close to the fronts. A rain cell above Poland blocked migration on the Eastern part of the domain. In contrast, a clear pathway is visible from Northern Germany through to Southwestern France.

While the estimation represents the most likely value of bird density at each node of the grid (e.g. Figure 5), a simulation generate several realizations (i.e. equiprobable outcomes of the migration process) that reproduce the space-time patterns of migration (e.g. Figure 6). As a consequence, only realizations are able to reproduce number during peak migration as noted when comparing the colorscale of Figure 5 and Figure 6. This becomes important when assessing for instance the total number of birds aloft.

**Figure 6.**
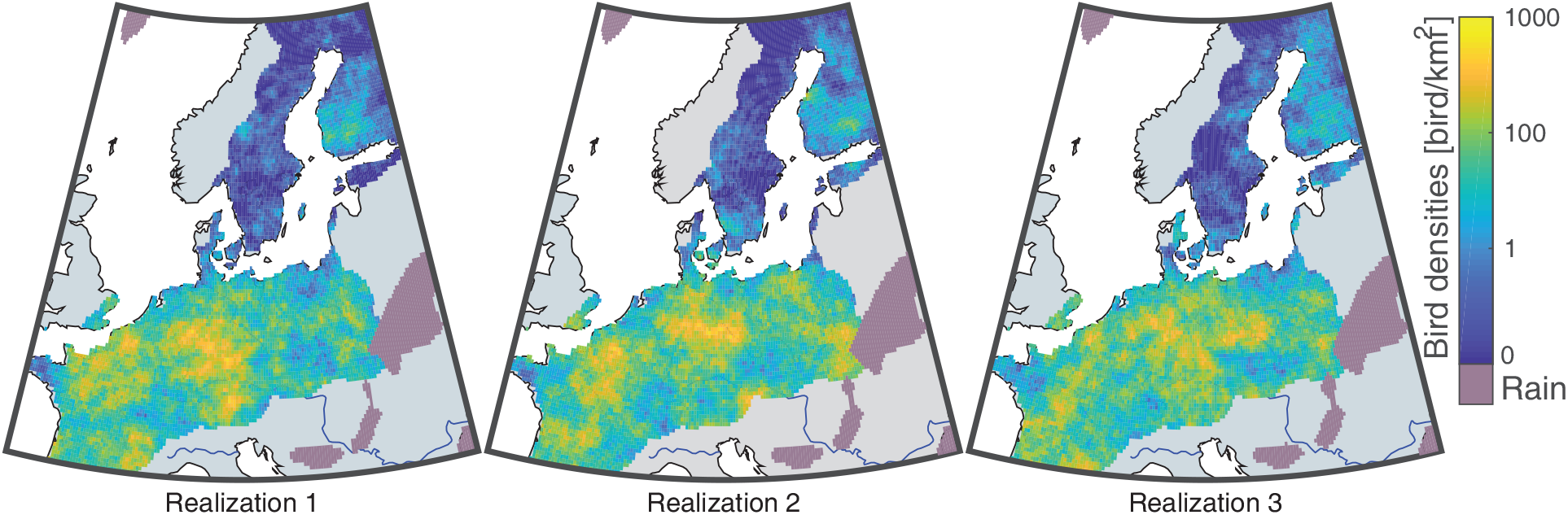
Snapshot of three different realizations showing peak migration (4 October 2016 21:30 UTC). The total number of birds in the air for these realisations was 119, 126 and 120 million respectively. Comparing the similarity and differences of bird densities patterns among the realisations illustrates the variability allowed by the stochastic model used. The texture of these realisations is more coherent with the observations than the smooth estimation map in Figure 5.

For each of the 100 realisations, we computed the total number of birds flying over the whole domain for each time step (Figure 7b). Within the time periods considered in this study, the peak migration occurred in the night of 4/5 October with up to 120 million [10-90 quantiles: 107-134] birds flying simultaneously. Computing this on sub-domains such as countries highlight the geographical differences in migration intensity. For instance, on the same night, France had 44 [39-51] million birds aloft (89 bird/km^2^), 20 [15-25] million for Poland (65 bird/km^2^), and only 8 [7-10] million in Finland (30 bird/km^2^) (Figure 7c).

**Figure 7.**
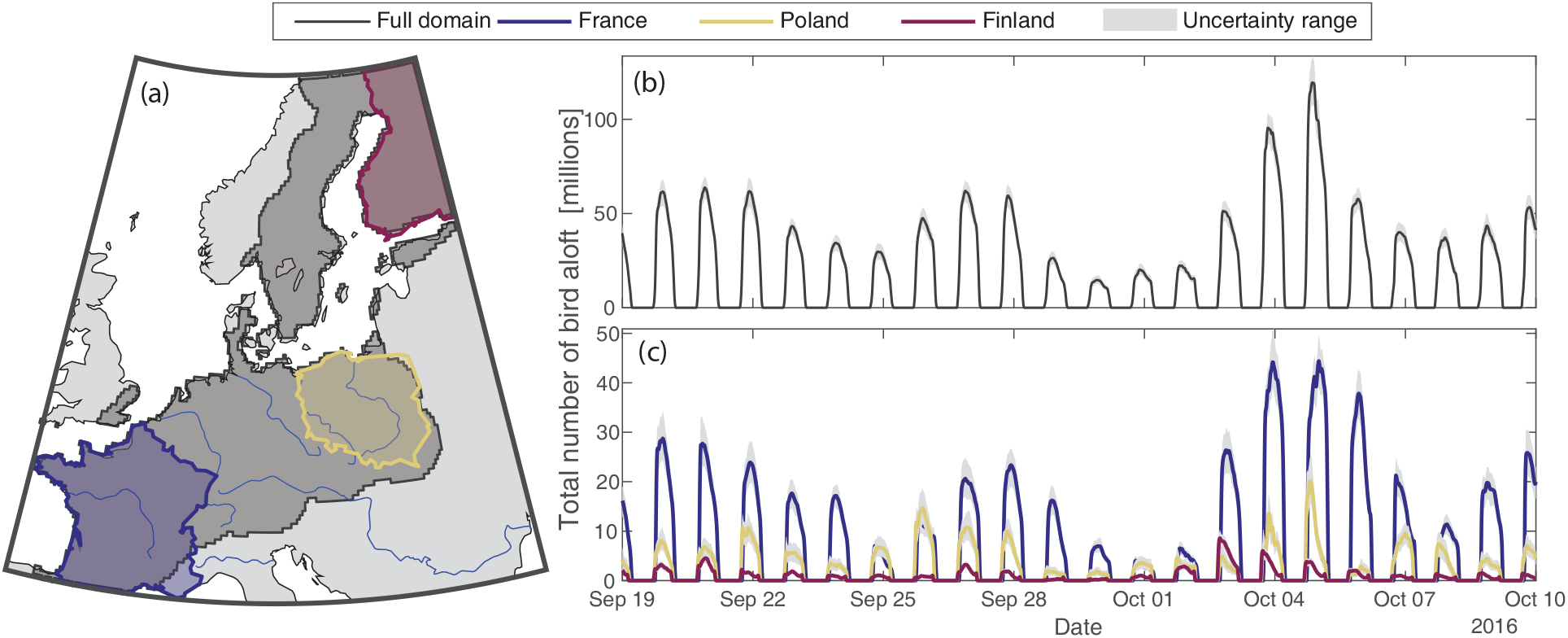
Averaged time series of the total number of birds in migration over the whole domain (black line), France (blue line), Poland (yellow line) and Finland (red line) and their associated uncertainty ranges (10-90 quantiles, light grey).

The spatio-temporal dynamics of bird migration can be visualized with an animated and interactive map (available online at birdmigrationmap.vogelwarte.ch with user manual provided in Supporting Information S5), produced with an open-source script (github.com/Rafnuss-PostDoc/BMM-web). In the web app, users can visualize the estimated maps or a single simulation maps animated in time, as well as time series of bird densities of any location on the map. In addition, it is also possible to compute the number of birds over a custom area and download all of these data through a dedicated API.

## Discussion

The model developed here can estimate bird migration intensity and its uncertainty range on a high-resolution space-time grid (0.2°lat. lon. and 15 min.). The highest total number of birds flying simultaneously over the study area is estimated to 120 million [10-90 quantiles: 107-134], corresponding to an average density of 52 birds/km^2^. This number illustrates the impressive magnitude of nocturnal bird migration, and resembles values of peak migration estimated over the USA with 500 million birds and a similar average density of 51 birds/km^2^ (Van Doren & Horton, 2018). For more local results, interactive maps of the resulting bird density are available on a website with a dedicated interface that facilitates the visualisation and the export of the estimated bird densities and their associated uncertainty (birdmigrationmap.vogelwarte.ch, see Supporting Information S5 for a user manual).

### Advantages and limitations

This paper presents the first spatio-temporal interpolation of nocturnal bird densities at the continental scale that accounts for sub-daily fluctuations and provides uncertainty ranges. In contrast to the methods based on covariates that are deemed more reliable for extrapolation in space and time (Erni et al., 2002; Van Belle, Shamoun-Baranes, Van Loon, & Bouten, 2007; Van Doren & Horton, 2018), our interpolation approach does not require any external covariate per se (e.g. temperature, rain, or wind). Although local features such as the approach of a rain front, the proximity to the ocean, or the presence of mountains will affect bird migration, these were not explicitly accounted for in the current model. However, their influence on bird migration is partially captured by the measurements of weather radars, so that, in turn, the interpolation implicitly accounts for them. Yet, if such covariates are available, the model can easily be adapted to incorporate information from these covariates through co-kriging (e.g. Chilés & Delfiner, 1999). However, adding these covariables to the interpolation is only advantageous if the correlation between bird densities and these covariates is stronger than the spatio-temporal correlation of the nearby radars.

In addition to quantifying bird migration at high resolution, we can also deduce the spatio-temporal scales at which migration is happening from the covariance function of the model (Figure S3-5 in Supporting Information S3). For instance, the amplitude of bird migration correlates over distances up to 700 km and over periods of up to 5.5 days (i.e. distance for which the auto-covariance has reached 10% of its baseline value, see Supporting Information S3 for details). These decorrelation ranges of the amplitude scale the spatio-temporal extent of broad front migration in the midst of the autumn migration season and highlight the importance of international cooperation for data acquisition and for spread of warning systems on peak migration events.

As a proof of concept, we used three weeks of bird densities data available on the ENRAM data repository (see Data Accessibility). As more data from weather radars become available, our analyses can be extended to year-round estimations of migration intensity at the continental scale, in Europe and in North America. We also importantly pre-processed the bird density data, i.e. restricted our model to nocturnal movements and applied a strict manual data cleaning. This is because the bird density data presently made available can be strongly contaminated with the presence of insects during the day, and birds flying at low altitude are not reliably recorded by radars because of ground clutter and the radar position in relation to its surrounding topography. Once the quality of the bird density data has improved, our model can be implemented in near-real-time and provide continuous information to the stakeholder, public and private sector.

Although we think that the model introduced here can already be a valuable tool (see below), we see several avenues for further development. For instance, in applications where the distribution of flight altitudes is crucial, the model can be extended to explicitly incorporate the vertical dimension. Furthermore, if fluxes of birds, i.e. migration traffic rates, are sought after, a similar geostatistical approach can be used to interpolate flight speeds and directions that are also derived from weather radar data.

### Applications

Many applied problems rely on high-resolution estimates of bird densities and migration intensities and the model developed here lays the groundwork for addressing these challenges. For instance, such migration maps can identify migration hotspots, i.e. areas through which many aerial migrants move, and thus, assist in prioritising conservation efforts. Furthermore, mitigating collision risks of birds by turning off artificial lights of tall buildings or shutting down wind energy installations requires information on when and where migration intensity peaks. The probability distribution function of our model can provide this as it estimates when and where migration intensity exceeds a given threshold. Such information can be used in shut-down on demand protocols for wind turbine operators, or trigger alarms to infrastructure managers.

## Acknowledgements

This study contains modified Copernicus Climate Change Service Information 2019. Neither the European Commission nor ECMWF is responsible for any use that may be made of the Copernicus Information or Data it contains. We acknowledge the European Operational Program for Exchange of Weather Radar Information (EUMETNET/OPERA) for providing access to European radar data, facilitated through a research-only license agreement between EUMETNET/OPERA members and ENRAM (European Network for Radar surveillance of Animal Movements). Mathieu Boos kindly provided the BirdScan data for Herzeele in France. We acknowledge the financial support from the Globam project funded by BioDIVERSA, including the Swiss National Science Foundation (31BD30 184120), Netherlands Organisation for Scientific Research (NWO E10008), Academy of Finland (aka 326315), BelSPO BR/185/A1/GloBAM-BE.

## Authors’ contributions

RN, LB, FL, BS conceived the study, RN, LB, GM designed the geostatistical model, RN developed and implement the computational framework, RN, LB, BS performed the analyses and wrote a first draft of the manuscript, with substantial contributions from all authors.

## Data Accessibility

- The Github page of the project (rafnuss-postdoc.github.io/BMM contains the MATLAB livescript for the interpolation rafnuss-postdoc.github.io/BMM/2016/html/Density inference cross validation.html) and the creation of the figures (rafnuss-postdoc.github.io/BMM/2016/html/paperfigure).
- Raw weather radar data are available on the ENRAM repository (github.com/enram/data-repository).
- The cleaned vertical time series profile are available on Zenodo (doi.org/10.5281/zenodo.3243397) (Nussbaumer, Benoit, Mariethoz, et al., 2019)
- The final interpolated maps are available on Zenodo (doi.org/10.5281/zenodo.3243466) (Nussbaumer, Benoit, & Schmid, 2019).
- The code of the website (HTML, Js, NodeJs, Css) are available on the Github page (github.com/rafnuss-postdoc/BMM-web)

## Supporting Information

### S1 Data pre-processing

Supporting Information S1 describes the full procedure applied to manually clean the raw time series of bird densities. The raw dataset have been previously published in (Nilsson et al., 2019) and is available on the ENRAM data repository (github.com/enram/data-repository). The steps detailed below are illustrated in Figure S1-1 for the radar located in Zaventem, Belgium (50°54’19”N, 4°27’28”E).

1. The data of 11 radars are discarded because their quality was deemed insufficient by visual inspection. The reasons for this poor quality are various: S-band radar type, altitude cut, poor processing or large gaps. The same radars were removed in (Nilsson et al., 2019). In addition, we also excluded radars of Bulgaria and Portugal (4 radars) from this study because of their geographic isolation and the necessity of spatial coherence in the methodology presented.
2. If rain is present at any altitude bin, the full vertical profile was discarded (blue rectangle in Figure S1-1). A dedicated MATLAB GUI was used to visualise the data and manually set bird densities to “not-a-number” in such cases.
3. Zones of high bird densities are sometimes incorrectly eliminated in the raw data (red rectangle in Figure S1-1). To address this, (Nilsson et al., 2019) excluded problematic time or height ranges from the data. Here, in order to keep as much data as possible, the data have been manually edited to replace erroneous data either with “not-a-number”, or by cubic interpolation using the dedicated MATLAB GUI.
4. Due to ground scattering, the lowest altitude layers are sometimes contaminated by errors, or excluded by the initial automatic cleaning procedure. This is solved by copying the first layer without error into to the lowest ones.
5. The vertical profiles are vertically integrated from the radar altitude (brown line in Figure S1-1c) and up to 5000 m a.s.l.
6. Finally, the data recorded during daytime are excluded. Daytime is defined at each radar by the civil dawn and dusk (sun 6°below horizon).

The resulting cleaned vertical-integrated time series of nocturnal bird densities are displayed in Figure S1-1d.

**Figure S1-1.**
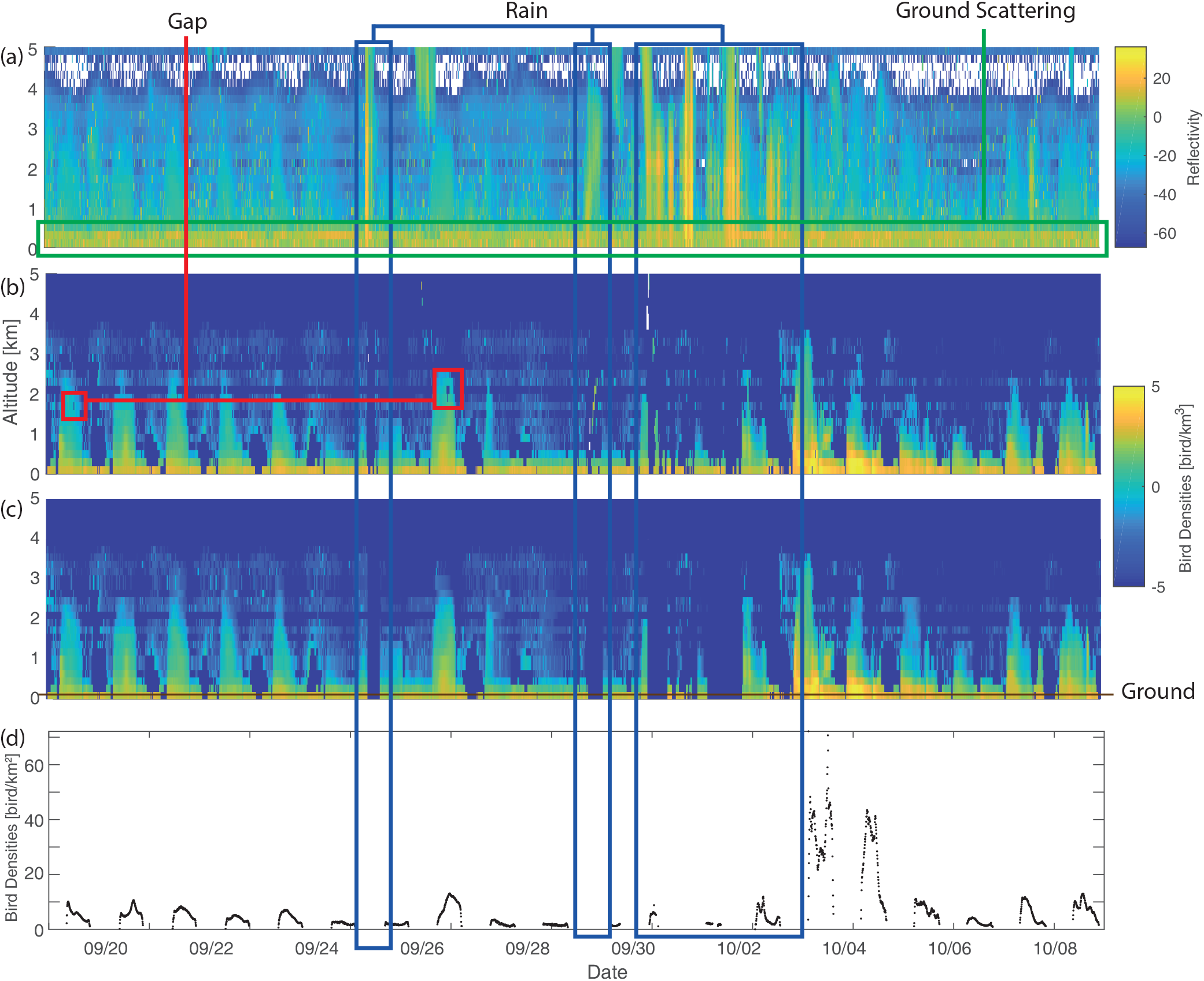
Example of the cleaning procedure from raw reflectivity to areal bird densities for the radar of Zaventem, Belgium (50°54’19”N, 4°27’28”E). (a) raw reflectivity measurements; (b) Automatically cleaned vertical bird profiles; (c) Manually cleaned vertical bird profiles; (d) Final bird densities (integrated over altitudes).

### S2 Details on the geostatistical model

Supporting Information S2 expends the explanation given in the section Geostatistical model. In particular, it provides the mathematical development for the kriging equations of the amplitude and residual as well as the reconstruction of the bird densities estimation.

#### Standardization

For the convenience of the kriging equation, we introduce the standardized (i.e. normal transformed) variable 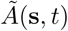 and 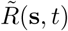 of the amplitude and residual respectively. Because the trend removed the average of the amplitude (i.e. **E**(*A*) = 0), the standardized amplitude is

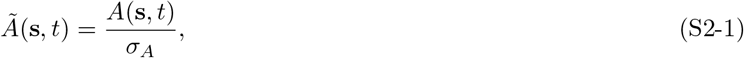

where 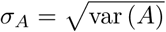 is the empirical standard deviation of *A*. The standardized residual is

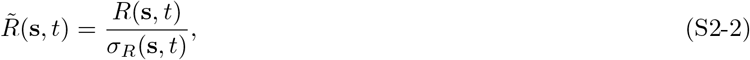

where the variance *σ*_*R*_(**s***, t*) is modelled by a polynomial function of the local *N N T* because of the strong correlation of the variance of the residual with the *N N T* (Figure S3-3 in Supporting Information S3),

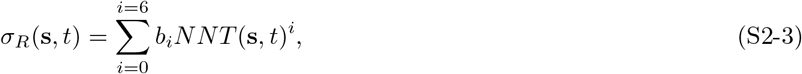

where *b*_*i*_ are the coefficients of the polynomial.

#### Covariance function

Both the amplitude and the residual are modelled as stationary processes which can be described with a covariance function (also called auto-covariance). Let us denote the generic standardized random variable 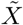 for either the standardized amplitude 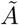 or the standardized residual 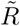. The covariance 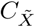 is a positive definite function depending only on the lag-distance (∆**s**, ∆*t*). In our model, we use covariance functions of Gneiting type (Gneiting, 2002)

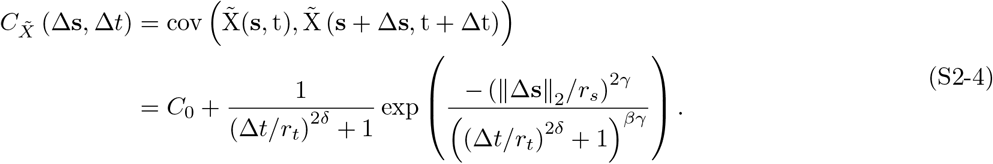

In this model, *r*_*t*_ and *r*_*s*_ are the scale parameters (in space and time respectively). They control the decorrelation distances and thus, the average extent and duration of the space-time patterns of 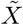. 0 *< δ*, *γ <* 1 are regularity parameters (in space and time respectively) and control the shape of the covariance function close to the origin. Values of *δ* and *γ* close to 0 lead to sharp variations at short lags, while values close to 1 lead to smooth variations of 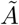. The separability parameter *β* controls the space-time interactions. When *β* = 0 the space-time interactions vanish and the covariance function becomes space-time separable. Finally, *C*_0_ is the nugget, which accounts for the uncorrelated variability of the process at hand.

#### Kriging

Both the standardized amplitude and residual can be estimated by kriging as explained below (Figure S2-1). Kriging provides an estimated value of the random variable 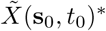 at the target point (**s**_0_*, t*_0_) based on a linear combination of observations at the *n*_0_ closest space-time locations 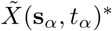 with

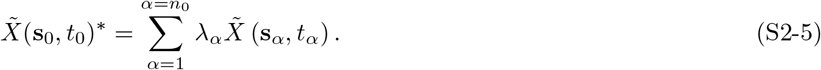

The kriging weights **Λ** = [*λ*_1_*, …, λ*_*n0*_]^*T*^ are derived from the covariance function of the random process 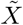. More precisely, the kriging weights are the solution of the following linear system, commonly referred to as the kriging system,

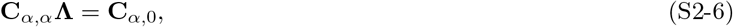

with **C**_*α,α*_ the covariance matrix between observations and **C**_*α*,0_ the covariance matrix between the observations and the target point. These covariances are computed using the fitted covariance function of Eqn. S2-4

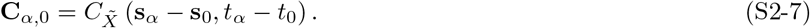

The kriging weights can be solved using 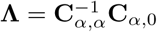, and used in Eqn. S2-5 to compute the kriging estimate.

In addition to the estimated value 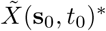, kriging also provides a measure of uncertainty with the variance of the estimation,

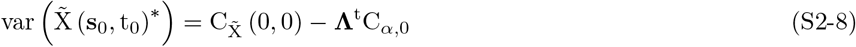

**Figure S2-1.**
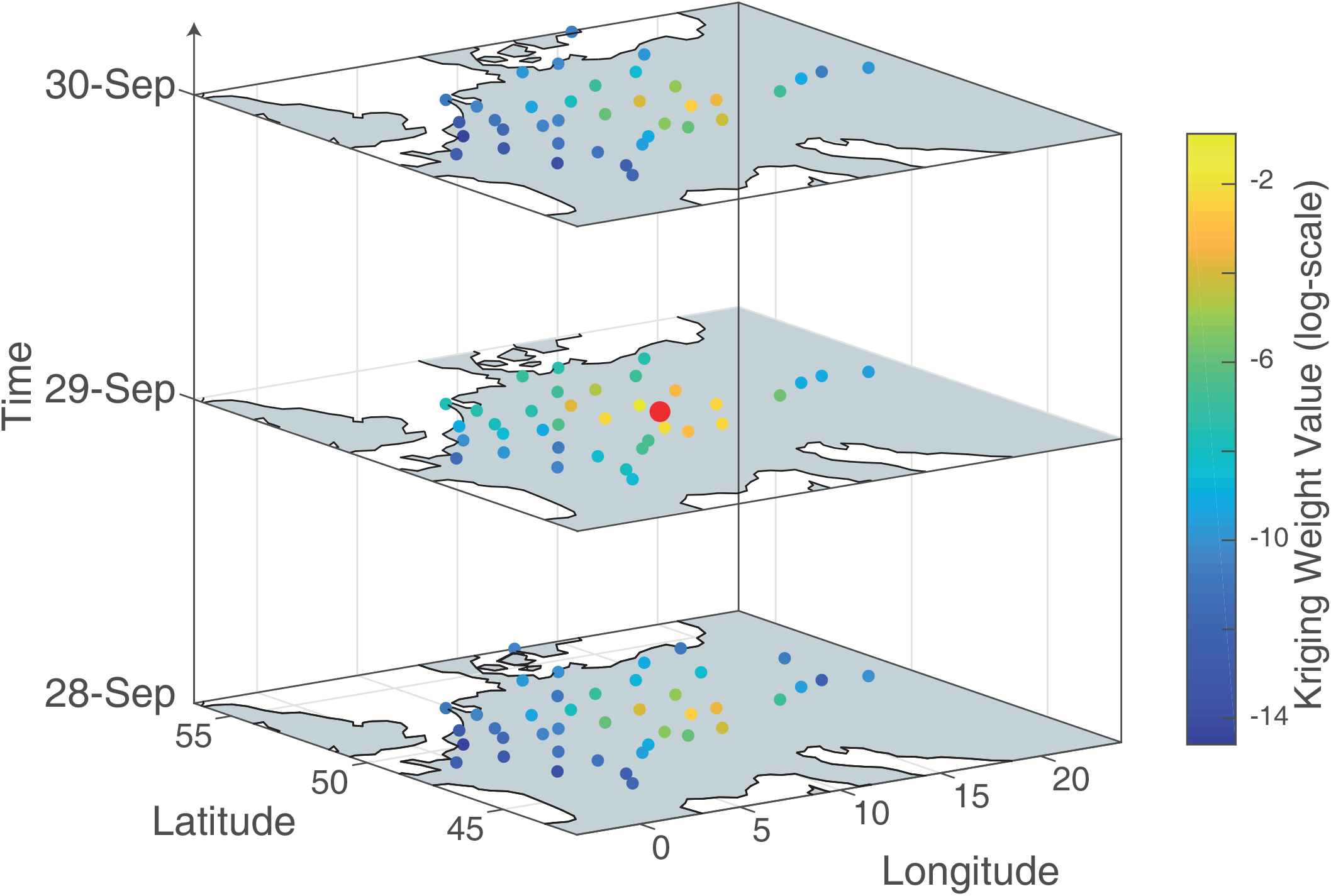
Illustration of the kriging weights computed for an estimation performed at the red dot location. Here we illustrate only the neighbours whithin +/- 1 day and a spatial neighbourhood of 600 km.

#### Reconstruction of the transformed bird densities

The transformed variable *Z* (**s**_0_*, t*_0_)^*p*^ is reconstructed by combining the deterministic parts (trend and curve) with the kriging estimation of the amplitude and residual as in Eqn. 1. Because *A* and *R* are normally distributed, *Z*^*p*^ is also normally distributed and its mean 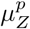 and variance 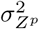 are sufficient to describe its distribution with

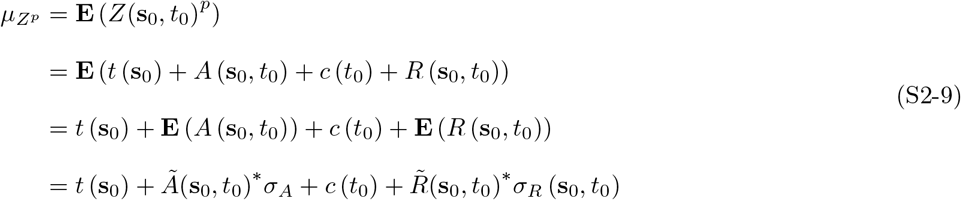

and as *A* and *R* are independent,

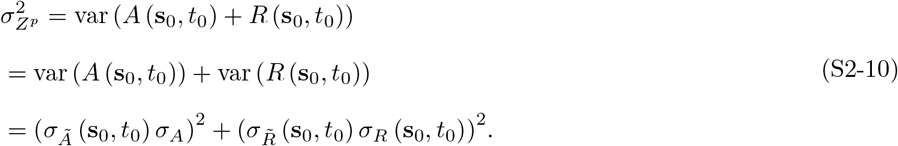

#### S2.1 Probability distribution function of bird densities

Because of the power-transform, the probability distribution function (pdf) of *Z*, denoted by *f*_*Z*_ (*z*), is non-trivial. However, the quantiles of this pdf can be derived analytically from the quantiles of the pdf of *Z*^*p*^ as detailed hereafter. We introduce the normally distributed variable *X* = *Z*^*p*^ with a pdf

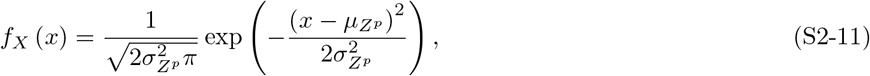

where *µ*_*Z*^*p*^_ and 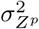 are the mean and variance of *Z*^*p*^ derived from Eqn. S2-9 and S2-10.

One possible way to compute the pdf of *Z*^*p*^ consists in computing the mean of several functions of this random variable. For any given measurable function *g*,

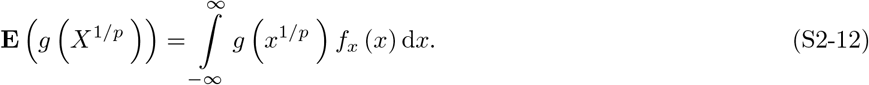

Using the change of variable *z* = *x*^1*/p*^, which leads to d*x* = *pZ*^*p−1*^*dz*, Eqn. S2-12 becomes

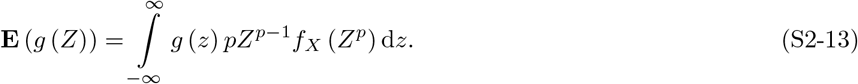

This equation allows identifying the pdf of *Z* as

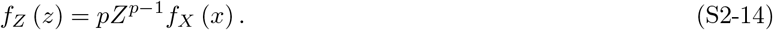

This last equation provides the analytical pdf of bird densities *Z*, as the pdf *f*_*X*_ (*x*) = *f*_*Z*^*p*^_ (*z*^*p*^) is fully known 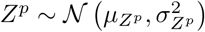.

### Quantile function

The probability distribution function of *Z* is non-symmetric and skewed, and therefore cannot be conveniently described with this expected value and variance. Instead, we use the quantile function *Q*_*Z*_ (*ρ*; **s**_0_*, t*_0_), which returns the bird density value *z* corresponding to a given quantile *ρ*

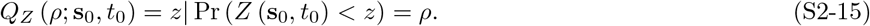

The quantile function allows to describe *Z* because the quantile value *ρ* is preserved through power transform. Therefore, the quantile function of *Z* is computed with

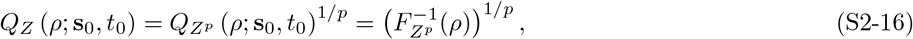

where *F*_*Z*^*p*^_ (*Z*^*p*^) is the cumulative distribution function of *Z*^*p*^(**s**_0_*, t*_0_).

### S3 Model parametrisation

Supporting Information S3 presents the method of parametrisation and discusses the meaning of model parameters in terms of bird migration.

#### Power transform

The value of power transformation *p* is inferred by maximizing the Kolmogorov-Smirnov criterion of the p-transformed observation data *Z*(**s***, t*)^*p*^. The Kolmogorov-Smirnov test (Massey, 1951) is testing the hypothesis that data *Z*(**s***, t*)^*p*^ are normally distributed. The optimal power transformation parameter was found for *p* = 1*/*7.4 and the resulting *Z*^*p*^ histogram is illustrated in Figure S3-1 together with the initial data *Z*.

**Figure S3-1.**
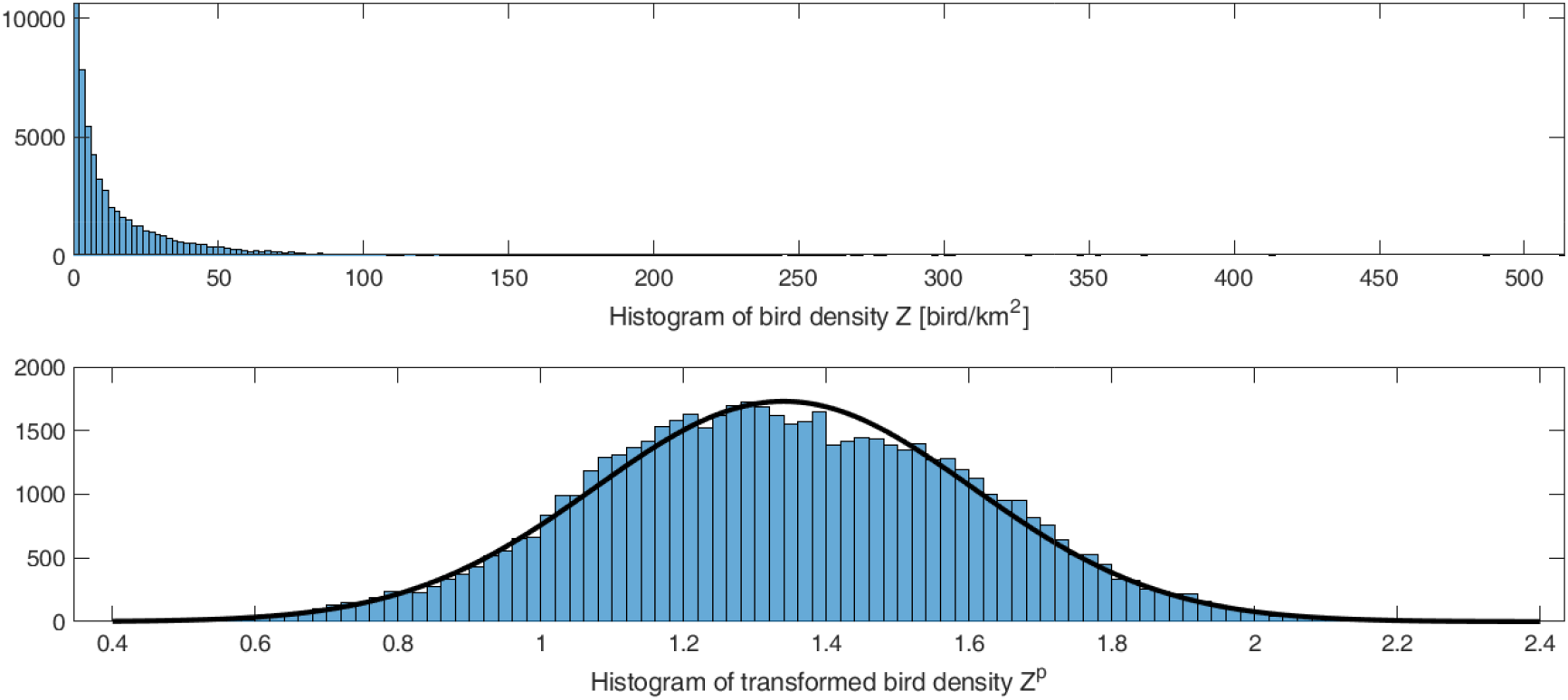
Histogram of the raw bird densities data *Z* (top) and the power transformed bird densities *Z*^*p*^ (bottom) for the calibrated parameter *p* = 1*/*7.4.

The fitted distribution shows that bird densities is highly skewed: the lowest 10% are below 1 bird/km^2^ while the upper 10% are above 50 bird/km^2^ with density up to 500 bird/km^2^. A power transformation on such skewed data creates significant non-linear effects in the back-transformation. For instance, the symmetric uncertainty of an estimated value in the transformed space (quantified by the variance of estimation) will become highly skewed in the original space. Consequently, the uncertainty of the estimation of bird densities is highly dependent on the value of the power transform: low densities estimations have a smaller uncertainty than high densities. This motivates the importance of providing the full distribution of the estimation. Indeed, the traditional central value (mean of 19 bird/km^2^ and median of 8 bird/km^2^) would be unable to capture the distribution adequately. Such effects also have consequences from an ecological/conservation point of view. Indeed, efficient protection of birds along the migration route (from artificial light or wind turbines) need to pay particular attention to the peak densities, where the majority of birds are moving in a few nights. These peaks can only be successfully reproduced by taking care of the high tail of the distribution. This is done here by using a full distribution for the estimation.

#### Trend and curve

The parameters of the deterministic components of the model (i.e. trend and curve) are calibrated based on the transformed data measured at the radar locations. This step involves the calibration of 3 parameters for the trend (*w*_lat_*, w*_lon_*, w*_0_), 6 for the curve (*a*_*i*_) and also the values of the amplitude for each radar and for each night (i.e. *n*_*radar*_ ⋅ *n*_*day*_ values). The calibration is performed by minimizing the misfit function between the modeled and the observed data. In practice, a local search algorithm is used (fminsearch function of MATLAB which uses a simplex algorithm). This local search algorithm requires initial values for each parameter, which are computed sequentially: (1) the initial trend is fitted to the average of the transformed bird densities for each radar, (2) the amplitudes are computed from the de-trended data, and (3) the parameters of the curve are derived by fitting a polynomial on the data corrected from the trend and amplitude effects. After convergence of the algorithm, the resulting misfit value becomes the value of the residual. The resulting planar trend is shown in Figure S3-2a together with the average transformed bird densities of each radar. The trend displays a strong North-South gradient, which can be explained by the larger migration activity in southern Europe during the study period. A 2-dimensional planar trend was initially tested in order to accommodate the northeast-southwest flyway. However, this more complex model did not significantly improved the fit to data, and has therefore been discarded. The de-trended values illustrated in Figure S3-2b are more stationary with the exception of Finland and Sweden. (Nilsson et al., 2019) also noted this difference between both countries, but excluded the fact that this difference is due to errors in the data since the southern Swedish radar shows consistent amounts of migratory movements with a neighbouring German site. Figure S3-2b highlights the central European continental flyway as illustrated by the arrow.

**Figure S3-2.**
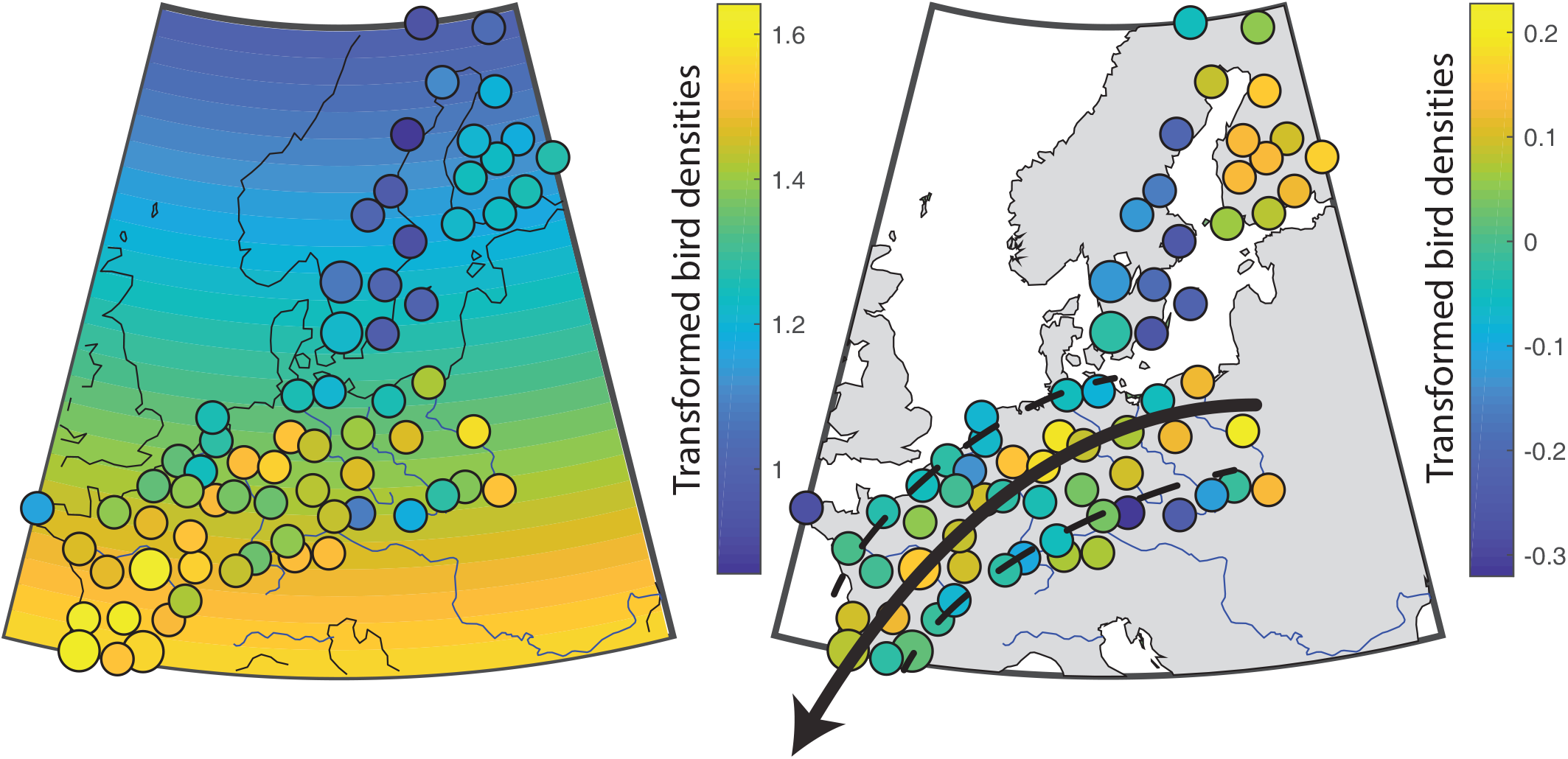
(left) Fitted trend with corresponding observation at radar location and (right) detrended data.

Figure S3-3 displays the fitted curve (black line) together with the calibration data. The curve reveals that the migration is mainly concentrated between 10-90% of the nighttime with larger densities of birds in the first half of the night. A slight asymmetry with steeper rise at the beginning of the night and smoother transition with the day is also visible. The larger variance of the data around the calibrated curve at the beginning and end of night is due to the power-transformation of the raw data and accounted for in the model. However, the large variance of the data used to adjust the curve model shows a significant intra-night variability in bird migration. This highlights the importance of modelling the intra-night fluctuations by the residual term and stresses the limitation of using nightly averages or single point observations (e.g. 3hr after sunset) if high precision estimates are required.

**Figure S3-3.**
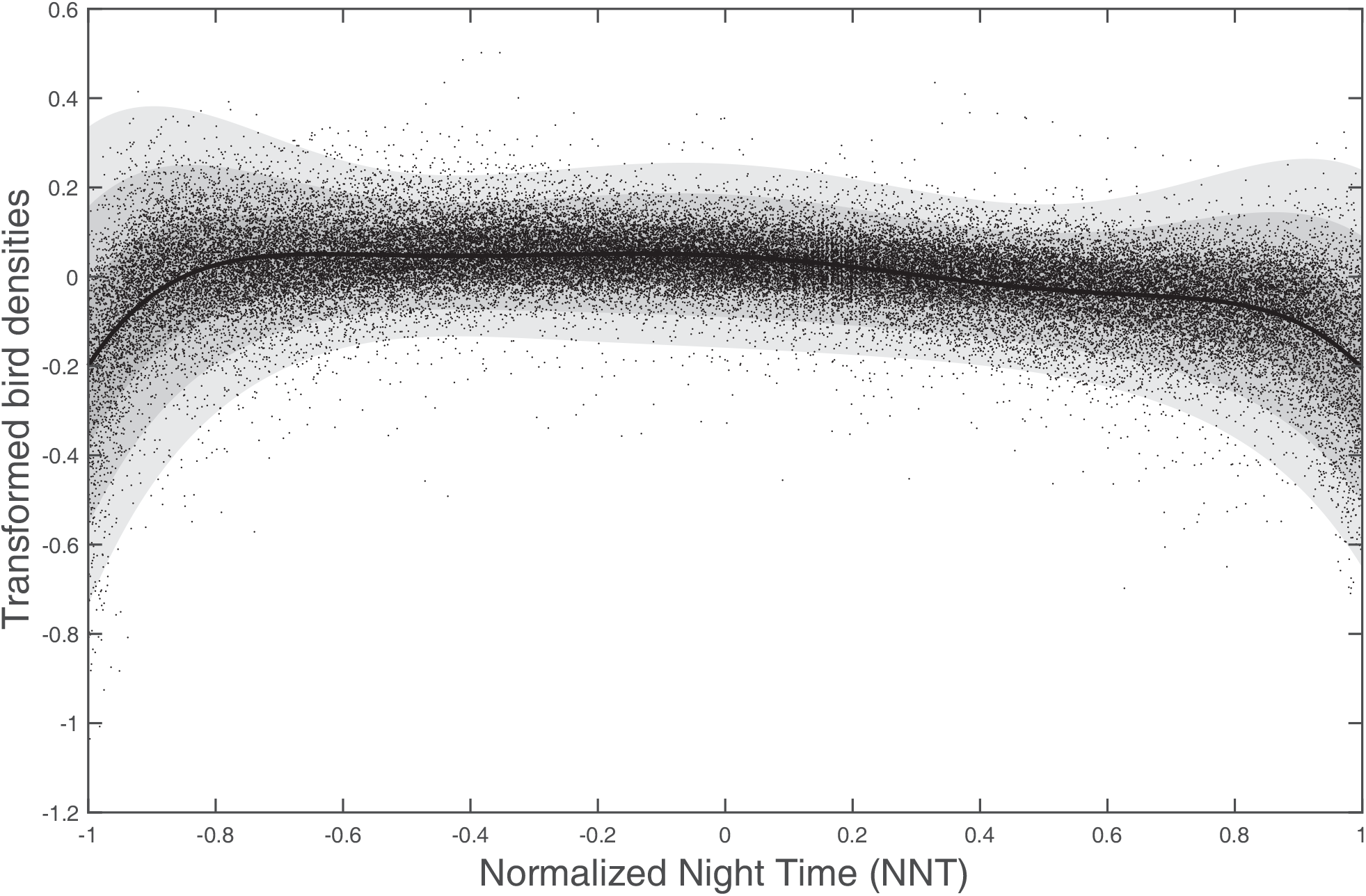
Individual observations (black dots) and adjusted model (black line) of the nightly curve. Shaded grey areas denote 1-, 2- and 3- sigma uncertainty ranges.

As no clear spatial patterns appear in the curve parameters, a single curve model is used for the entire study domain. The suitability of this unique curve model is validated by the spatially-uncorrelated mean and variance of the residual signal displayed in Figure S3-4.

**Figure S3-4.**
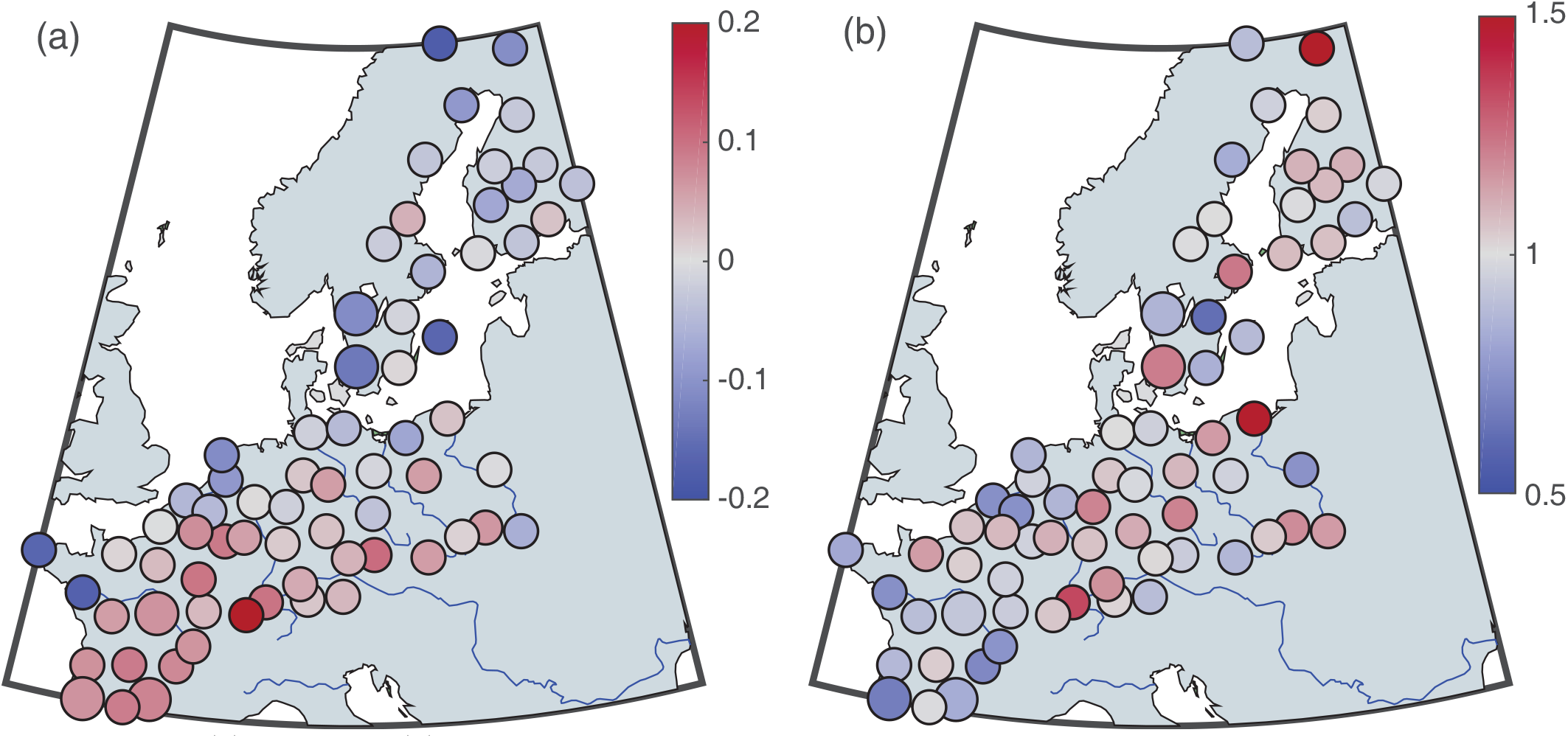
(a) Mean and (b) standard deviation of residual value for each radars.

### Amplitude and residual

The parameters of the space-time covariance function (*n, r_t_, r_s_, δ, γ, β*) of the amplitude and residual are inferred from empirical covariances derived for several lag-distances and lag-times on an irregular grid. Then, the parameters of the Gneiting covariance function are inferred by minimizing the root mean square error between empirical and modelled covariances. Both covariance functions of the amplitude and residual are best fitted with a separable model (*β* = 0), meaning that they can be fully described by the product of a spatial covariance function (Figure S3-5a and c) and a temporal covariance function (Figure S3-5 b and d).

### Covariance function of the amplitude and residual

The calibrated covariance functions provide information about the degree of spatial and temporal correlation of the bird migration process. The spatial covariance of the amplitude (Figure S3-5a) shows that the nocturnal bird densities are well-correlated for locations separated by less than 500 km, and completely uncorrelated for more than 1500 km. The temporal covariance has an asymptotic behavior and never decreases under 0.2 (Figure S3-5b). This non-zero sill can be due to either a remaining temporal non-stationarity in the dataset, or systematic errors in radar observations (caused by e.g. different types of technology or local topography affecting migration). Note that since the covariance is evaluated only on a discrete 1-day lag-distance, the shape of the covariance between 0 and 1 is artificially created to fit the Gneiting function. Overall, the temporal correlation of the amplitude is weak with only 40% for the covariance for lag-1. It is important to recall that since the weather radars are relatively well-spread (Figure 1b), the spatial covariance of both the amplitude and residual is poorly constrained for lag-distances below 100 km, and consequently the importance of the nugget (Eqn. S2-4) is unknown. The temporal correlation of the residual is very high for short lags (*<*2 hr) and indicates consistent measurements of each weather radars. The small covariance value for larger lag-distances demonstrates that the curve account for most of the stationary component at this scale.

**Figure S3-5.**
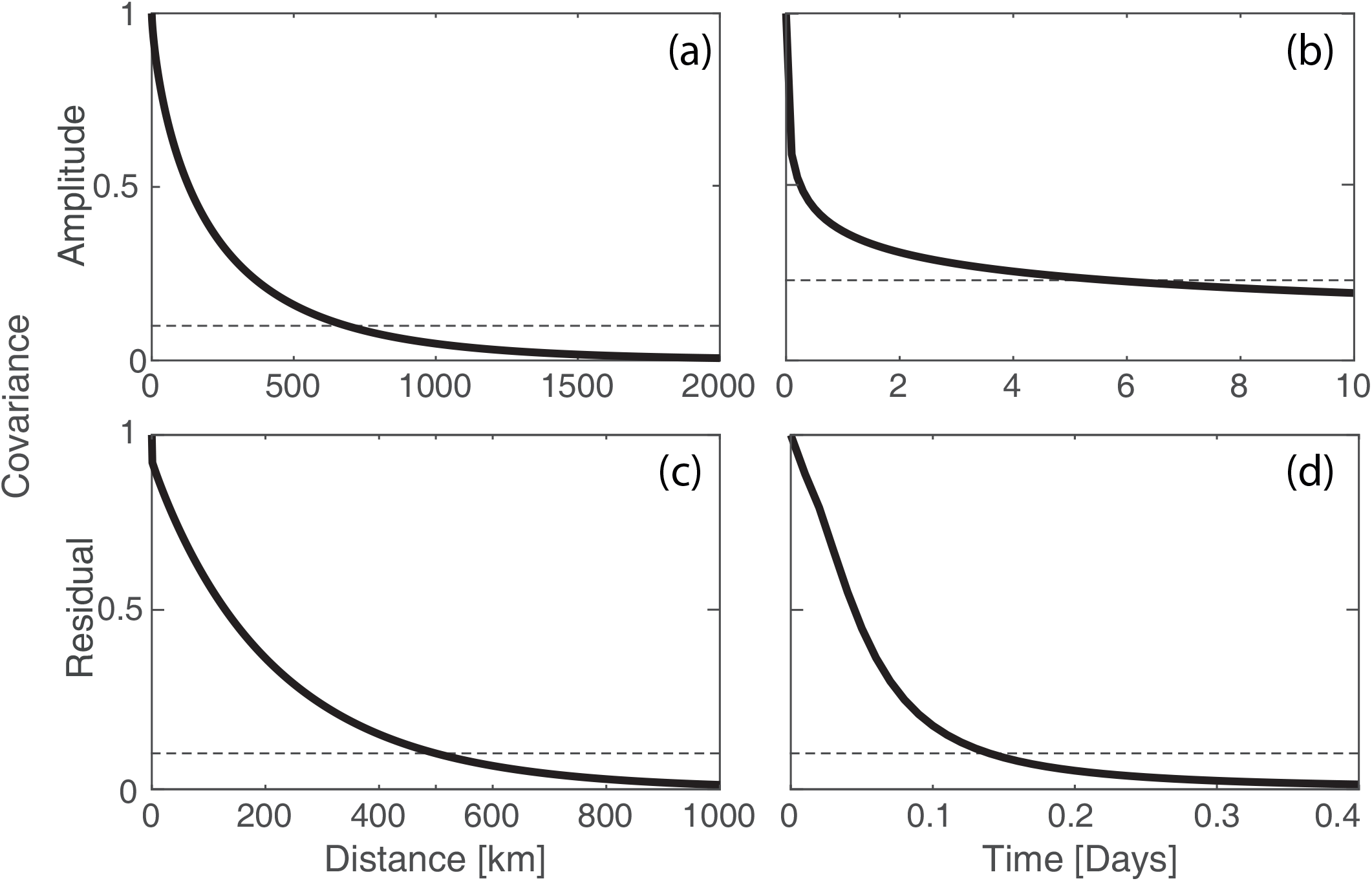
Illustration of the calibrated covariance function of (a-b) amplitude and (c-d) residual.

The table below summarizes the fitted parameters.

**Table 1.** Calibrated parameters

|  |  |
| --- | --- |
| Power Transformation | $p = 0.135$ |
| Trend | $w_0 = 2.62$ , $w_{lat} = -0.024$ |
| Covariance of amplitude | $nugget/sill = 0\%$, $\beta = 0$, $r_t = 0.39$, $r_s = 174$, $\delta = 0.18$, $\gamma = 0.43$ |
| Curve | $a = [-0.720.120.76 - 0.05 - 0.29 - 0.070.05]$ |
| Curve variance | $b = [-0.010.030.00 - 0.010.000.00]$ |
| Covariance of residual | $nugget/sill = 7.7\%$, $\beta = 0$, $r_t = 0.048$, $r_s = 206$, $\delta = 1$, $\gamma = 0.47$ |

### S4 Cross-validation

Supporting Information S4 extends the results of the Cross-validation section. First of all, Figure S4-1 displays the histogram of the normalized errors of kriging (Eqn. 5) when all data across all radars are combined. The mean of the distribution is close to zero which indicates that the estimation is unbiased (i.e. in average, the estimation is neither underestimating (mean below 1) nor overestimating (mean above 1)). However, its variance is below 1, which indicates a slight overestimation of the uncertainty range (i.e. in average, the uncertainty ranges are too wide).

**Figure S4-1.**
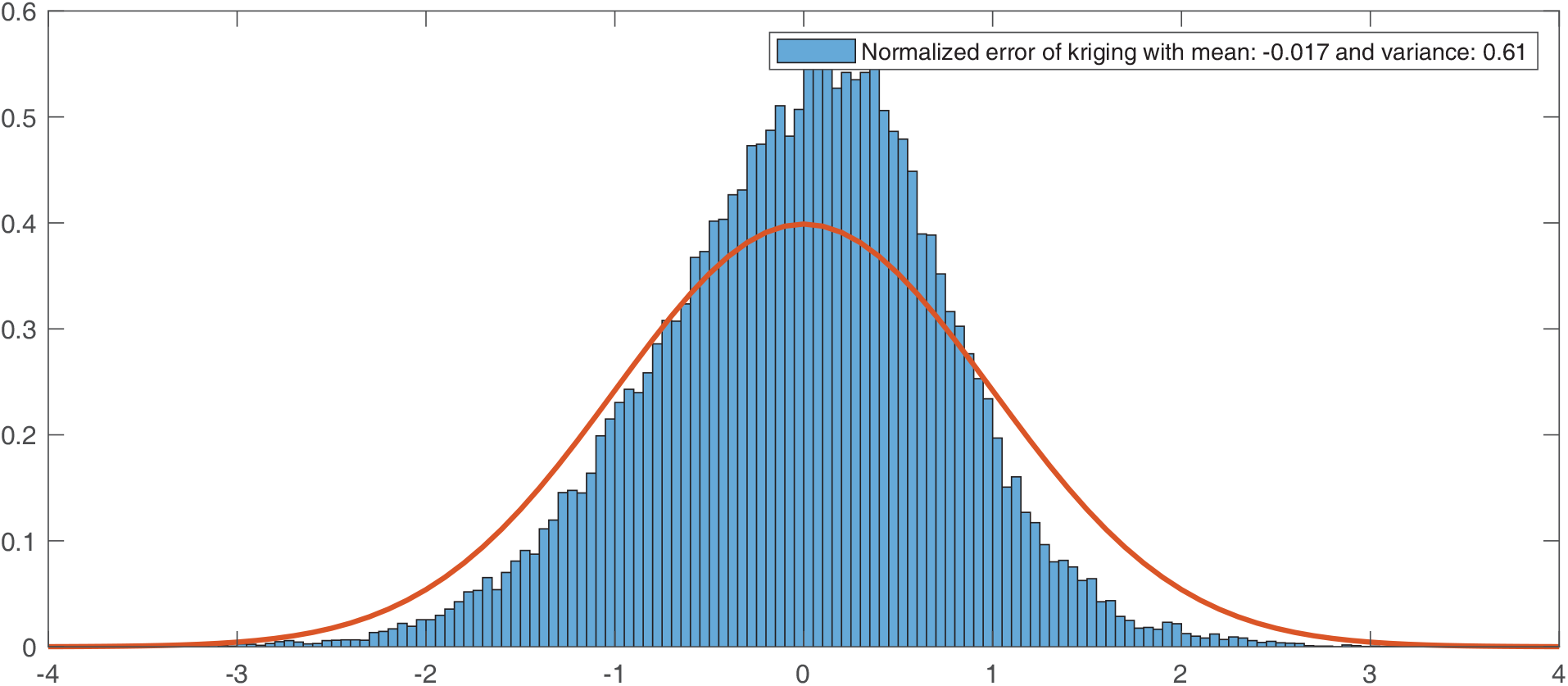
Histogram of the normalized error of kriging for all radars combined. The red curve is the standard normal distribution which should be matched by the histogram.

Next, the normalized kriging error is assessed for each radar (Figure S4-2). The resulting distributions indicate that the goodness of the estimation is different for each radar. In Figure S4-3, their means and standard deviations do not reveal any spatial pattern, thus suggesting no spatial biases.

**Figure S4-2.**
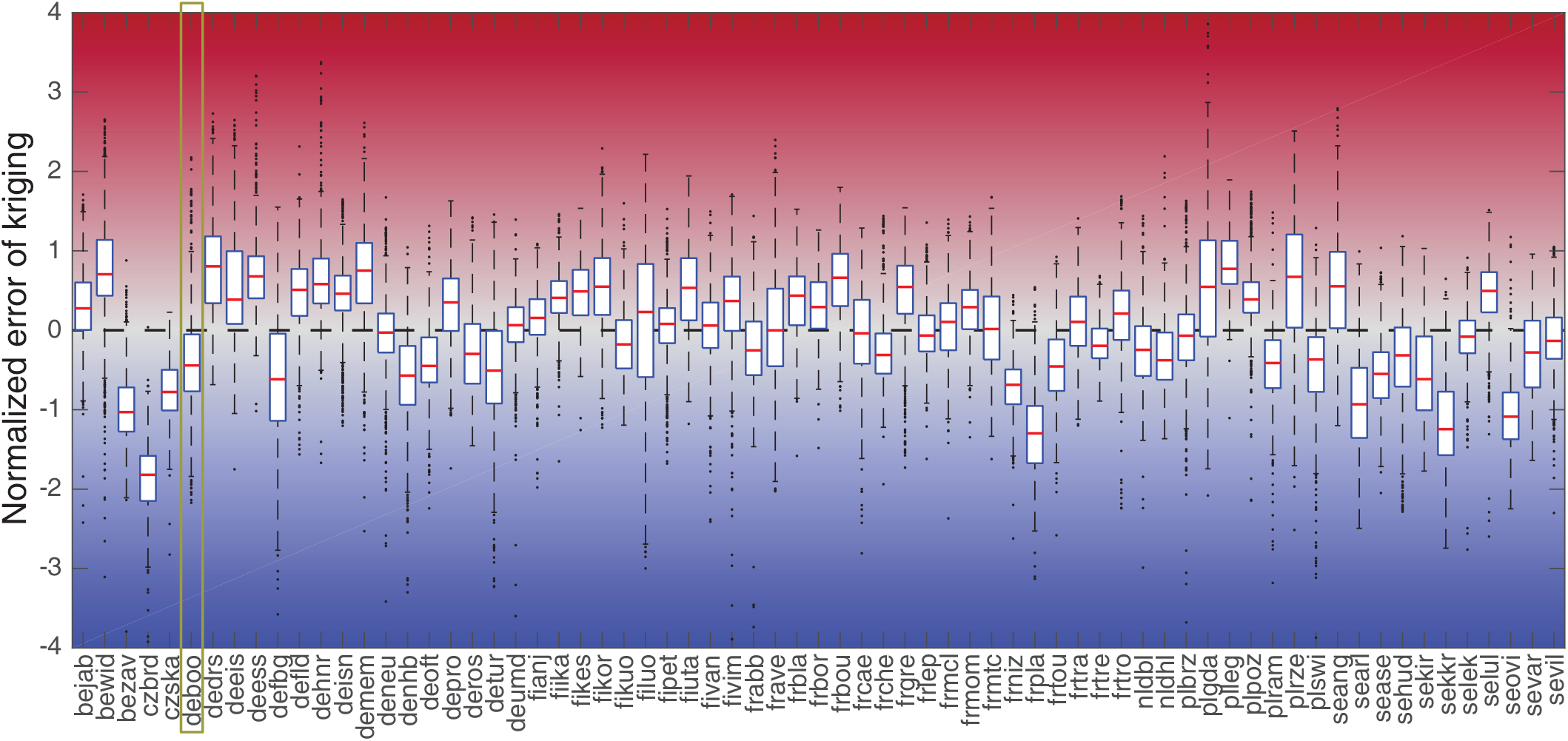
Boxplot of the normalized kriging error for each radar. A negative (positive) value indicates an underestimation (overestimation) of the method. The cross-validation for the Czech (czbrd, czska) and Swedish radars (se***) shows a constant underestimation (except for selul). Overall, the goodness of the estimation is variable and radar-dependent.

**Figure S4-3.**
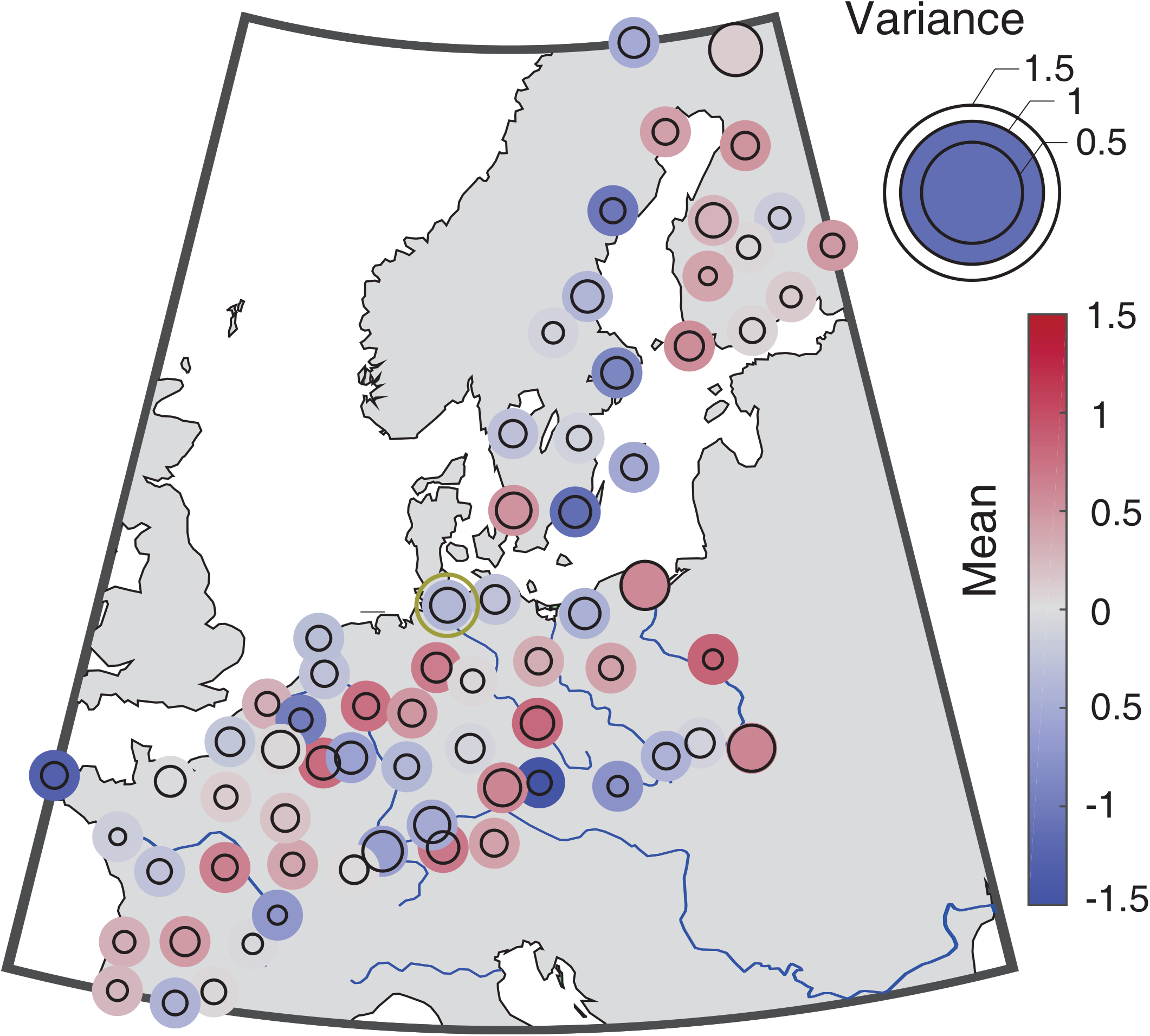
Mapping of the mean and variance of the normalized error of kriging for each radar. The reproduction of the variance is illustrated by a black circle, for which a perfect variance would match the colour circle and a smaller circle indicates an under-confidence (uncertainty range too large).

The cross-validation is further illustrated in Figure S4-4 for a specific radar located in Boostedt, Germany (54°00’16”N, 10°02’49”E) indicated with gold circle in Figure S4-3. For this radar, the general pattern of the signal is well estimated for both the nightly amplitude and the intra-night variation. Bird densities are underestimated during the peak migration occurring at 4th and 5th of October, is but the estimated value remains within the uncertainty bounds.

**Figure S4-4.**
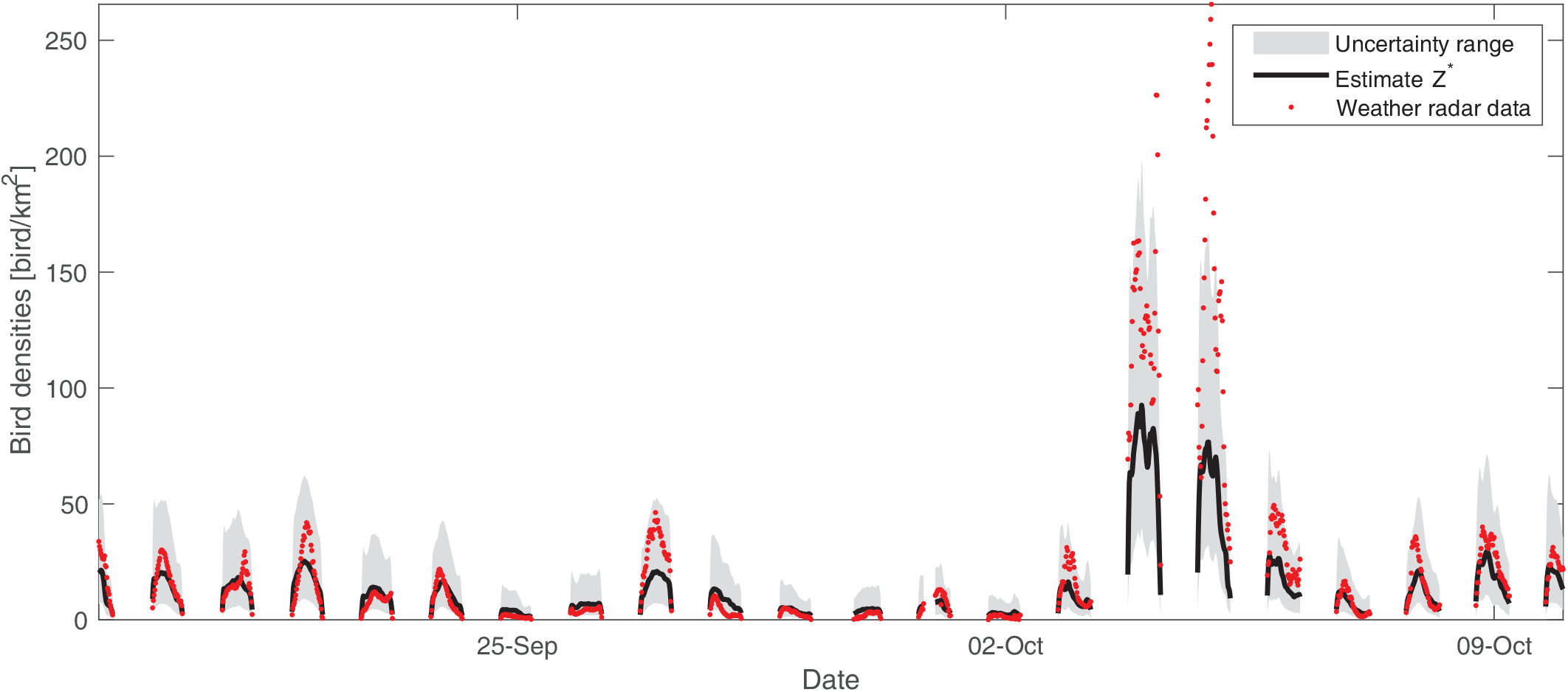
Comparison of bird densities estimated with the model in a cross-validation setup and observed by the weather radar for the radar located in Boostedt, Germany (54°00’16”N, 10°02’49”E). The uncertainty range is defined as the 10 and 90 quantiles.

### S5 Manual for website interface

The following Supporting Information describes the web interface developed for visualization and querying of the interpolated data. The website is available at birdmigrationmap.vogelwarte.ch and the code at github.com/Rafnuss-PostDoc/BMM-web. Figure S5-1 displays the web interface along with the possible interactions, which are further detailed below.

**Figure S5-1.**
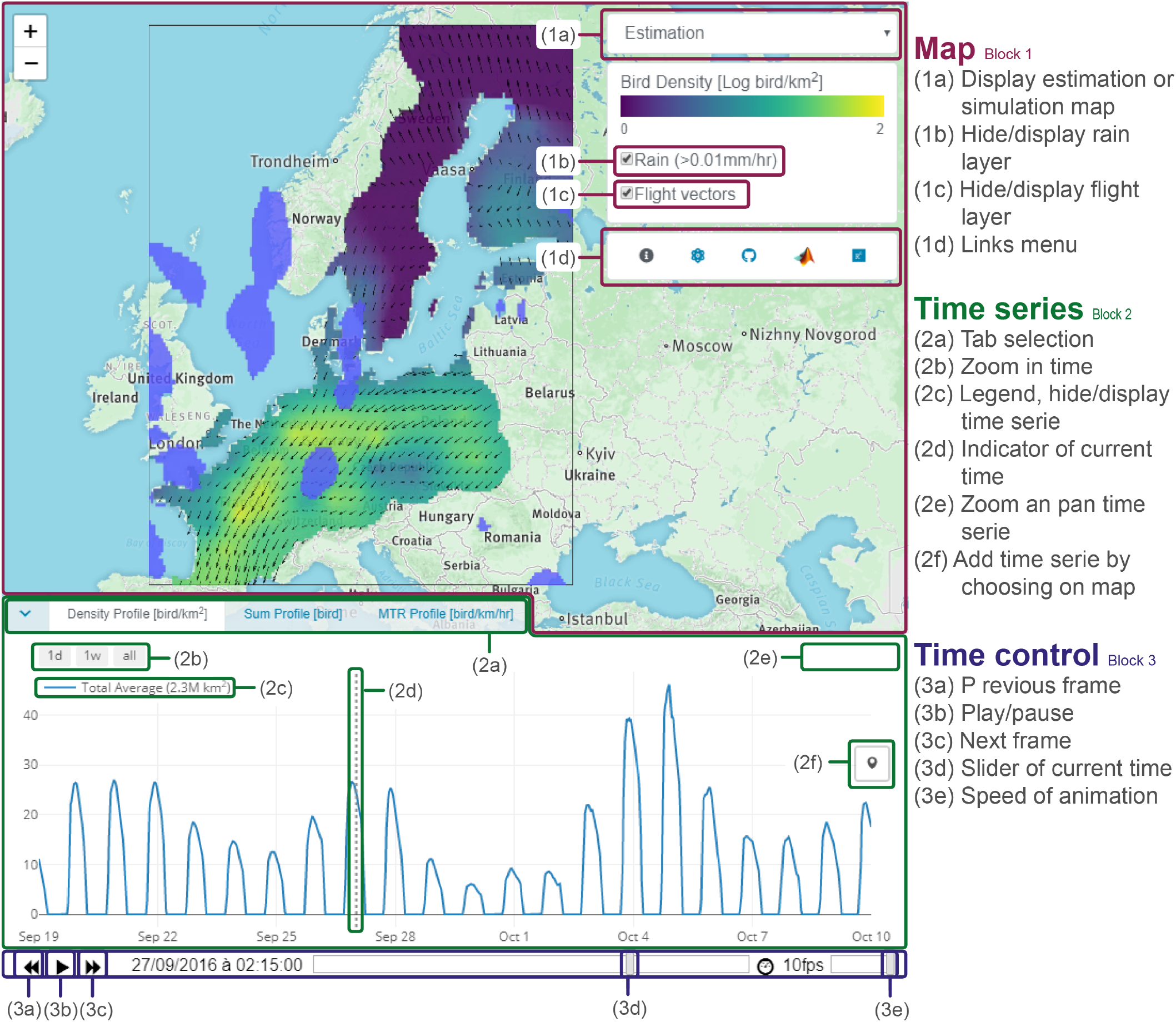
Website interface with identification key for the interactive components of each block

### Block 1: interactive map

The main block of the website is a map with a standard interactive visualization allowing for zoom and pan. On top of this map, three layers can be displayed:

- Layer 1 corresponds to bird densities displayed in a log-color scale. This layer can display either the estimation map, or a single simulation map by using the drop-down menu (1a).
- Layer 2 corresponds to the rain (rainy areas are in light blue), which can be hidden/displayed with a checkbox (1b)
- Layer 3 corresponds to the bird flight speed and direction, displayed by black arrows. The checkbox allows to display/hide this layer (1c). The last item on the top-right menu is the link menu (1d).

### Block 2: time series

The second block (hidden by default on the website) shows three time series, each in a different tab (2a):

- Densities profile shows the bird densities [bird/km^2^] at a specific location.
- Sum profile shows the total number of bird [bird] over an area.
- MRT profile shows the mean traffic rate (MTR) [bird/km/hr] perpendicular to a transect.

A dotted vertical line (2d) appears on each time series to show the current time frame displayed in the map. Basic interactive tools for time series include zooming on a specific time period (day, week or all period) (2b) and general zoom and auto-scale (2e). Each time series can be hidden and displayed by clicking on its legend (2c). The main feature of this block is the ability to visualise bird densities time series for any location chosen on the map. For the densities profile tab, the button with a marker icon (2f) lets you plot a marker on the map, and displays the bird densities profile with uncertainty (quantile 10 and 90) on the time series corresponding to this location. You can plot several markers to compare the different locations (Figure S5-2). Similarly, for the sum profile, the button with a polygon icon (2f) lets you draw any polygon and returns the time series of the total number of birds flying over this area. For the MTR tab, the flux of birds is computed on a segment (line of two points) by multiplying along the segment the bird densities with the local flight speed perpendicular to that segment.

### Block 3: time control

The third block shows the time progression of the animated map with a draggable slider (3d). You can control the time with the buttons play/pause (3b), previous (3a) and next frame (3c). The speed of animation can be changed with a slider (3e).

### API

An API based on mangodb and NodeJS is available to download any of the time series described in Block 2. Instructions can be found at github.com/Rafnuss-PostDoc/BMM-web#how-to-use-the-api

### Examples

**Figure S5-2.**
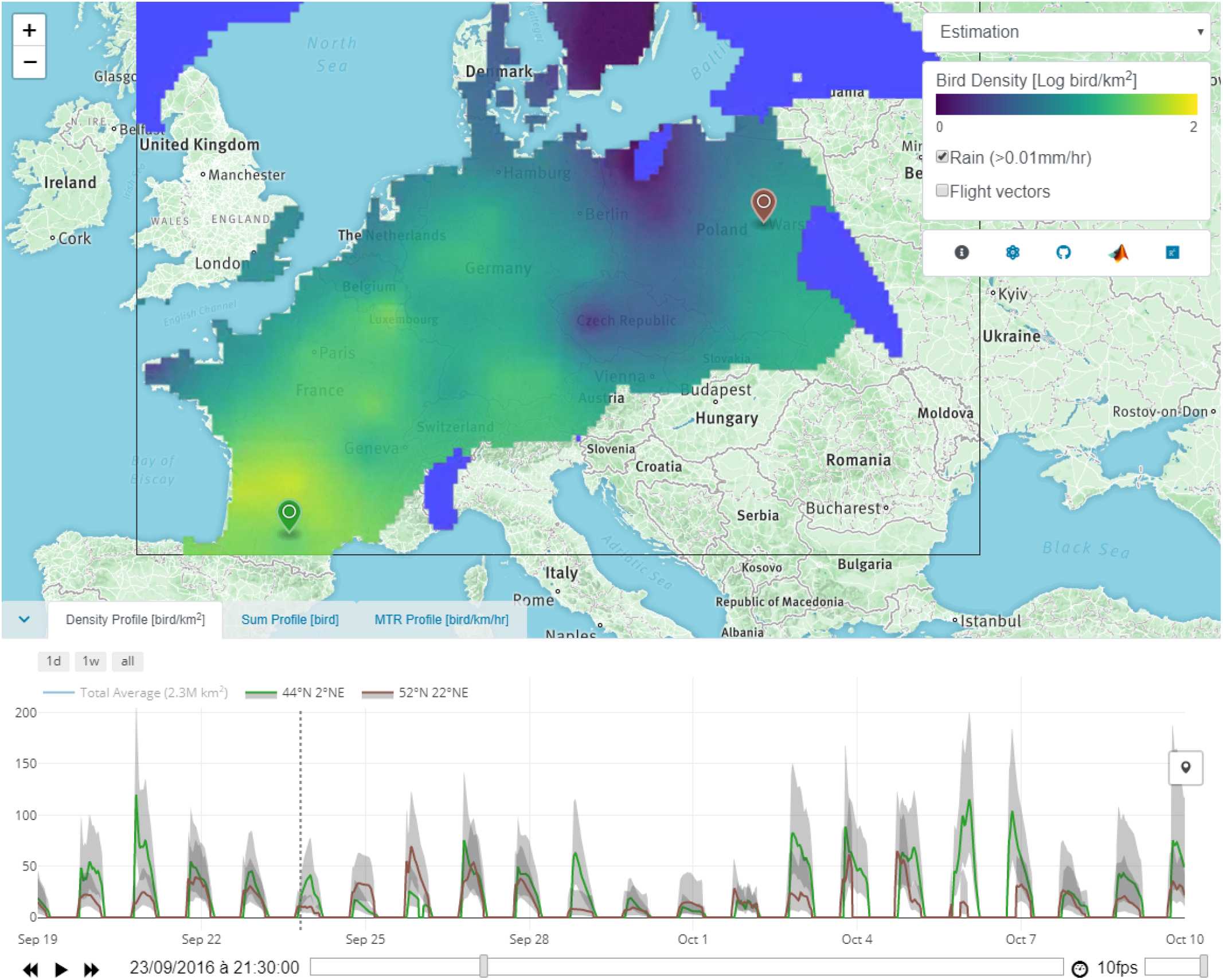
Print-screen of the web interface developed to visualize the dataset. The map shows the estimated bird densities for the 23rd of September 2016 at 21:30 with the rain mask appearing in light blue on top. The domain extent is illustrated by a black box. The time series in the bottom shows the bird densities with quantile 10 and 90 at the two locations symbolized by the markers with corresponding color on the map. The button with a marker symbol on the right side of the time series allows to query any location on the map, and to display the corresponding time series.

**Figure S5-3.**
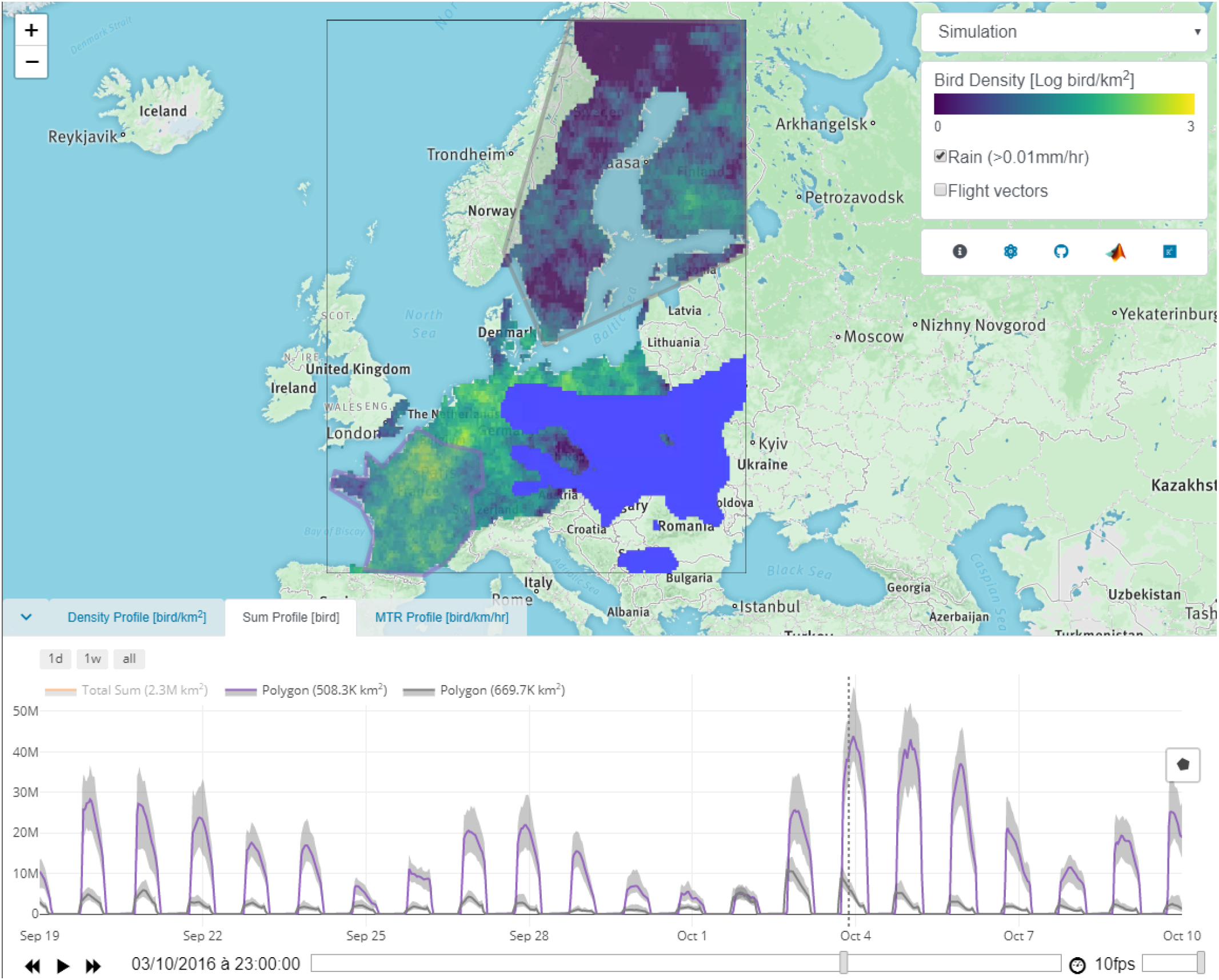
Print-screen of the web interface with the simulation map for the 3rd of October at 23:00. The bottom panel shows the time series of the total number of birds corresponding to the polygon drawn over the map according to their colour. The button with the polygon symbol on the right side of the time series allows to query the total number of birds flying any polygon drawn on the map.

## Notes

https://rafnuss-postdoc.github.io/BMM/

https://zenodo.org/record/3243397

https://zenodo.org/record/3243466

